# GABA-mediated inhibition in visual feedback neurons fine-tunes *Drosophila* male courtship

**DOI:** 10.1101/2023.01.25.525544

**Authors:** Yuta Mabuchi, Xinyue Cui, Lily Xie, Haein Kim, Tianxing Jiang, Nilay Yapici

**Author notes:** Correspondence: Nilay Yapici.

## Abstract

Vision is critical for the regulation of mating behaviors in many species. Here, we discovered that the *Drosophila* ortholog of human GABA_A_-receptor-associated protein (GABARAP) is required to fine-tune male courtship by modulating the activity of visual feedback neurons, lamina tangential cells (Lat). GABARAP is a ubiquitin-like protein that regulates cell-surface levels of GABA_A_ receptors. Knocking down *GABARAP* or *GABA*_*A*_ *receptors* in Lat neurons or hyperactivating them induces male courtship toward other males. Inhibiting Lat neurons, on the other hand, delays copulation by impairing the ability of males to follow females. Remarkably, the human ortholog of *Drosophila* GABARAP restores function in Lat neurons. Using *in vivo* two-photon imaging and optogenetics, we show that Lat neurons are functionally connected to neural circuits that mediate visually-guided courtship pursuits in males. Our work reveals a novel physiological role for GABARAP in fine-tuning the activity of a visual circuit that tracks a mating partner during courtship.

## Introduction

In many species, vision is used in a variety of fundamental behaviors such as navigation, reproduction, aggression, and prey hunting. Vision is especially critical in regulating social behaviors as visual cues can determine whether to approach or avoid a conspecific during social interactions. For example, the male peacock spider (*Maratus volans*) uses species-specific visual displays to communicate its reproductive capacity to females and attract their attention (Girard et al., 2011). Zebrafish (*Danio rerio*) evaluate visual features and biological motion to decide whether to interact with conspecifics. (Nunes et al., 2020). Paper wasps (*Polistes fuscatus*) rely on visual features to discriminate their nest mates (Sheehan and Tibbetts, 2011). Although the role of vision in regulating social behaviors has been well established in behavioral ecology, the molecular and neural mechanisms that evaluate and process complex visual information during social behaviors remain poorly understood in any species.

The fly, *Drosophila melanogaster*, uses vision to regulate a variety of its behaviors, including locomotion (Tammero and Dickinson, 2002; Triphan et al., 2010), navigation (Dewar et al., 2015; Srinivasan et al., 1999), courtship (Cook, 1979, 1980; Krstic et al., 2009; Markow, 1987), and learning (Neuser et al., 2008; Ofstad et al., 2011). As in vertebrates, the fly visual system is organized in anatomically segregated neuropils devoted to visual information processing: lamina, medulla, lobula, and lobula plate (Behnia and Desplan, 2015; Davis et al., 2020; Fischbach and Dittrich, 1989; Sanes and Zipursky, 2010). From the early visual processing areas, the lamina and the medulla, distinct classes of visual output neurons carry motion and color vision information to different central brain regions via a diverse population of visual projection neurons (Wu et al., 2016; Yamaguchi et al., 2008). Distinct classes of visual projection neurons regulate various behavioral responses in flies. For example, the lobula columnar (LC) neurons project from the lobula and/or lobulate plate to the central brain and respond to specific visual features and motions (Aptekar et al., 2015; Keleş and Frye, 2017; Klapoetke et al., 2022; Morimoto et al., 2020; Ribeiro et al., 2018; Städele et al., 2020; Wu et al., 2016). In addition to visual projection neurons, in the fly optic lobe, multiple classes of visual feedback neurons carry signals from the central brain or other regions of the optic lobes to the medulla and the lamina (Tuthill et al., 2013; Tuthill et al., 2014). The functions of most visual feedback neurons are poorly understood.

Vision also plays a critical role in *Drosophila melanogaster* courtship (Connolly et al., 1969; Cook, 1979, 1980; Markow, 1987; Markow and Manning, 1980; Ribeiro et al., 2018). During courtship behavior, male flies display a complex repertoire of actions to stimulate female flies to copulate with them. These include orienting toward a target female, chasing the female, tapping the female abdomen, ipsilaterally extending wings and producing a courtship song, licking female genitalia, and bending the abdomen to copulate (Bastock and Manning, 1955; Dickson, 2008; Greenspan and Ferveur, 2000; Hall, 1994; Yamamoto and Koganezawa, 2013). While male flies rely on chemosensation to assess the species, sex, and receptivity of suitable females and to initiate courtship (Clowney et al., 2015; Dweck et al., 2015; Fan et al., 2013; Kallman et al., 2015; Kurtovic et al., 2007; Thistle et al., 2012), they use vision for locating and tracking the female’s movements. Male flies with impaired vision take longer to chase and successfully copulate with females (Connolly et al., 1969; Cook, 1979, 1980; Krstic et al., 2009; Markow, 1987; Markow and Manning, 1980; Ribeiro et al., 2018). The delay in copulation in visually impaired flies is caused mainly by the male’s inability to track the female’s movements during courtship (Cook, 1979, 1980; Ning et al., 2022; Ribeiro et al., 2018). Recently, a class of visual projection neurons, lobula columnar cell type 10a (LC10a), has been shown to respond to small moving objects and regulate male tracking behavior during courtship. Inhibiting the activity of LC10a neurons in males impairs their ability to orient toward or maintain proximity to females (Hindmarsh Sten et al., 2021; Ribeiro et al., 2018).

Although vision is critical for the mating success of *Drosophila* males, visual cues alone are insufficient to trigger the full repertoire of courtship actions (Agrawal et al., 2014; Kohatsu et al., 2011; Pan et al., 2012). For example, the activity of LC10a visual projection neurons is enhanced when males are sexually aroused, and LC10a can initiate robust courtship only in sexually aroused males (Hindmarsh Sten et al., 2021; Ribeiro et al., 2018). In the fly brain, sexual arousal is regulated mainly by a class of male-specific neurons that express the sex-determination factor fruitless (fruM) (Manoli et al., 2005; Stockinger et al., 2005). A small population of fruitless positive neurons in the posterior fly brain (Fru-P1) can generate a persistent arousal state and stimulate male courtship when activated (Jung et al., 2020; Kimura et al., 2008; Kohatsu et al., 2011; Von Philipsborn et al., 2011). How sexual arousal regulates LC10a activity or, in general, visual processing during courtship is poorly understood.

Here, we identified a GABA_A_-receptor-associated protein (GABARAP) as a regulator of male courtship behavior in *Drosophila*. GABARAP (also known as Atg8a) belongs to a family of ubiquitin-like proteins that regulate multiple intracellular processes, including autophagy (Martens and Fracchiolla, 2020; Mizushima, 2020; Shpilka et al., 2011). In the canonical autophagy pathway, GABARAP (Atg8a) works together with other Atg proteins to form an autophagosome (Mizushima and Komatsu, 2011; Nakatogawa et al., 2007; Noda et al., 2013; Shpilka et al., 2011). In mammals, Atg8 family proteins are subdivided into two classes: the GABA type A receptor-associated protein (GABARAP) subfamily and the microtubule-associated protein one light chain 3 (LC3) subfamily (Jacquet et al., 2021; Jatana et al., 2020; Shpilka et al., 2011). Although these sub-families are initially thought to play redundant roles in autophagy, recent studies suggest they also have autophagy-independent functions (Schaaf et al., 2016; Shpilka et al., 2011). For example, mammalian GABARAPs regulate neuronal excitability by modulating the membrane trafficking and synaptic localization of GABA_A_ receptors (Kanematsu et al., 2007; Leil et al., 2004; Ye et al., 2021). Previous studies in flies have focused on the autophagy-related functions of GABARAP (Atg8a) (Bali and Shravage, 2017; Jipa et al., 2021; Ratliff et al., 2015; Simonsen et al., 2008). However, our study discovered that GABARAP (Atg8a) has autophagy-independent functions in the fly visual system. We found that neuronal knockdown of *GABARAP* or *GABA*_*A*_ *receptors* in a small population of visual feedback neurons, lamina tangential cells (Lat), elevated male courtship toward other males. Lat-induced male-male courtship defect is sensory context-dependent and relies on light conditions. Loss of GABARAP (Atg8a) function in Lat neurons did not impair autophagy; instead, we demonstrated that GABARAP regulates male courtship by modulating the activity of Lat neurons. We further showed that *Drosophila* GABARAP and human GABARAP share remarkable sequence identity, and *Drosophila* GABARAP function in Lat neurons can be rescued by its human ortholog. Finally, we found that Lat neurons regulate male courtship via a neural circuit that controls visually guided courtship pursuits. Our work revealed a novel physiological role for *Drosophila* GABARAP in regulating visual processing during male courtship. The dysfunctions in GABA_A_ receptor signaling have been associated with many psychiatric disorders that impact social behaviors, including autism spectrum disorder and bipolar disorder (Fatemi et al., 2002; Fatemi et al., 2009; Wellcome Trust Case Control, 2007). Here, we showed that GABARAP/GABA_A_ receptor signaling modulates the activity of neural circuits that regulate social behaviors in flies. We speculate that our study might provide insights into how GABA_A_ receptor signaling in the brain modulates social behaviors in flies and other animals and may help us understand the molecular and neural mechanisms underlying social deficits seen in human psychiatric disorders.

## Results

### GABARAP is required in the male nervous system to suppress courtship toward other males

We identified *Atg8a* (hereafter named *GABARAP*) as a regulator of male courtship behavior in a genome-wide transgenic RNA interference (RNAi) screen for genes required in the nervous system for reproduction (Bussell et al., 2014). RNAi-mediated knockdown of *GABARAP* in male flies using a pan-neuronal driver, *elav-GAL4* (*elav>*), induced male courtship toward other males (Figures 1A and S1A). *Elav>GABARAP-RNAi* males showed robust courtship actions toward target males without showing signs of motor deficits or general hyperactivity (Video S1). Male-to-female courtship was normal in *elav>GABARAP-RNAi* males (Figure 1B). Next, to investigate whether Atg proteins that are suggested to interact with GABARAP (Atg8a) (Mizushima, 2020; Nakatogawa et al., 2007) are also required to suppress male-male courtship, we knocked them down in the nervous system. *Elav>Atg3* males were lethal. Most of the *elav>Atg-RNAi* males tested (except for *Atg4a* and *Atg10*) courted target males (Figure S1B). Our results revealed that the functions of multiple *Atg* genes, including *GABARAP* (*Atg8a*), are required in the nervous system to regulate male courtship, specifically to suppress male courtship toward other males.

**Figure 1.**
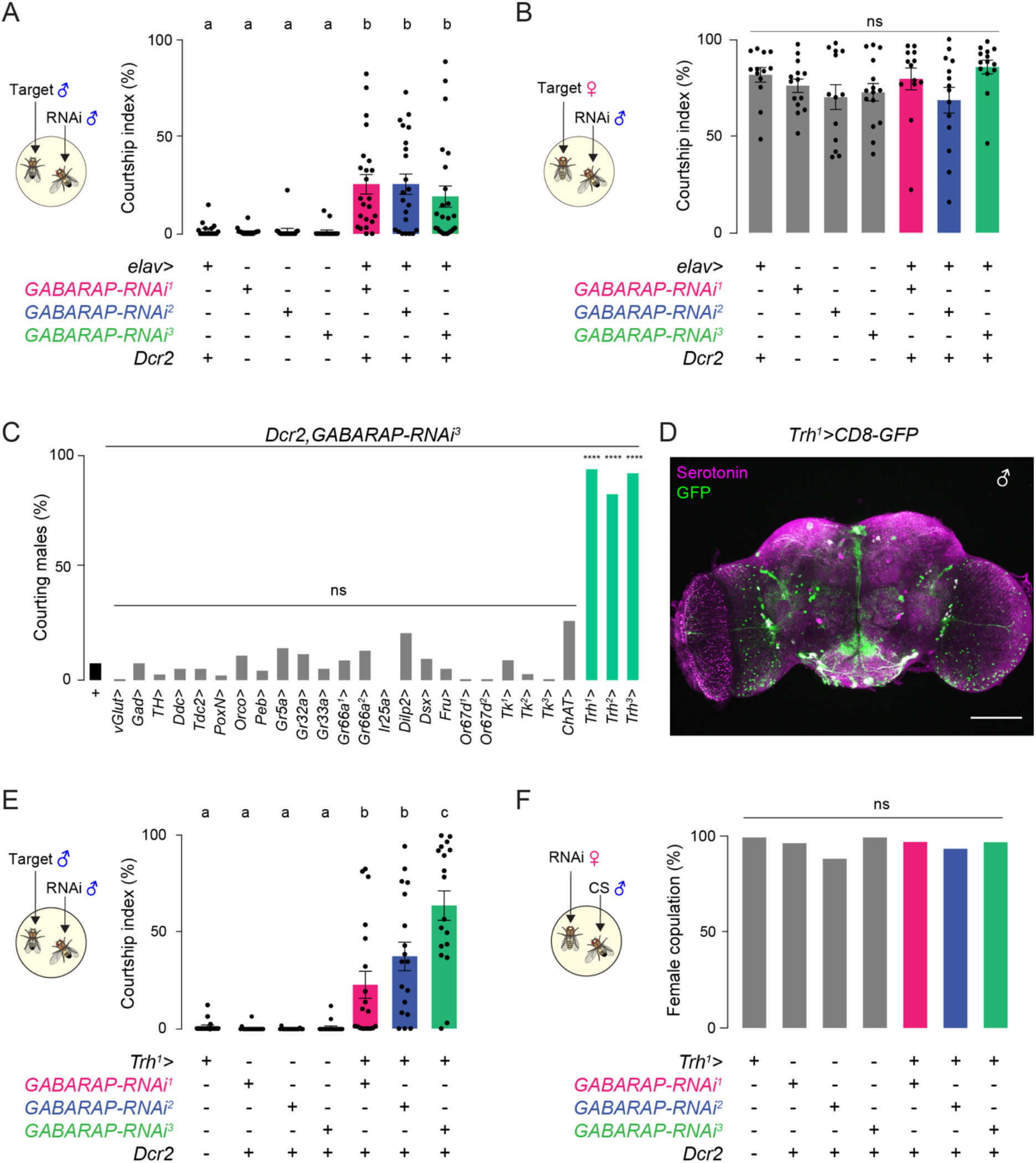
*GABARAP* is required to suppress male courtship toward other males. (A) Courtship quantification of *elav>GABARAP-RNAi* males and controls toward a male target (One-way ANOVA with Tukey’s test, p<0.05, mean ± SEM, n=18-24 flies). (B) Courtship quantification of *elav>GABARAP-RNAi* males and controls toward a female target (One-way ANOVA with Tukey’s test, ns, mean ± SEM, n=13-15 flies). (C) Percentage of males courting a male target in various GAL4s crossed to *GABARAP-RNAi*^*3*^. Positive GAL4s are colored in green (Fisher’s exact test, ****, p<0.0001; n=17-56 flies). (D) *Trh*^*1*^*>CD8-GFP* brain stained with anti-GFP (green) and anti-serotonin (magenta) (scale bar=50μm). (E) Courtship quantification of *Trh*^*1*^*>GABARAP-RNAi* and control males toward male targets (One-way ANOVA with Tukey’s test, p<0.05, mean ± SEM, n=18-19 flies). (F) Female receptivity quantification of *Trh*^*1*^*>GABARAP-RNAi* and control females as percent copulated in 20min (Fisher’s exact test, ns, n=34-80 flies).

### GABARAP is required in visual feedback neurons to suppress male courtship toward other males

To identify neurons in which *GABARAP* regulates male courtship, we carried out cell-type specific knockdown in molecularly defined populations of neurons related to courtship regulation, sensory perception, and neuromodulation and checked for male courtship defects. This allowed us to pinpoint GABARAP function to a population of neurons expressing the *tryptophan hydroxylase* gene (*Trh*) (Coleman and Neckameyer, 2005) (Figures 1C and 1D). *Trh>GABARAP-RNAi* males courted target males, displaying the full repertoire of male courtship actions (Figures 1E, S1C, and S1D, Video S2). We did not find any obvious courtship defects in *Trh>GABARAP-RNAi* females; they mated with target males at similar levels to controls (Figure 1F).

*Trh* gene encodes the rate-limiting enzyme in serotonin biosynthesis (Coleman and Neckameyer, 2005). *Trh>* labels most of the serotonergic neurons in the fly brain (n=∼90 per hemisphere). These neurons are classified into different clusters based on the location of their cell bodies (Figures 2A and 2B) (Alekseyenko et al., 2010; Pooryasin and Fiala, 2015; Sitaraman et al., 2012; Vallés and White, 1988). To identify a smaller subset of serotonergic neurons in which GABARAP is required to suppress male-male courtship, we expressed *GABARAP-RNAi* in different serotonergic clusters. We found that knocking down *GABARAP* in the lateral protocerebrum (LP) cluster was sufficient to induce male courtship toward other males (Figures 2C-2F). We further narrowed down the subset of neurons within the LP cluster using the split-GAL4 binary expression system (Luan et al., 2006; Pfeiffer et al., 2010). This intersectional genetic strategy allowed us to restrict GABARAP function to a small population of serotonergic visual feedback neurons (lamina tangential cells, Lat, n=∼6 per hemisphere) (Nässel, 1988, 1991) (Figures 2G and S2A-S2C). In all split-GAL4 drivers tested (*Lat*^*1*^*>, Lat*^*2*^*>, Lat*^*3*^*>*), *Lat>GABARAP-RNAi* males courted other males significantly more than genetic controls (Figures 2H and S2D, Video S3). The male-male courtship defect was not caused by changes in Lat neurogenesis, as we did not observe differences in the number of Lat neurons (Figure S2G) or gross anatomical changes in Lat wiring between control and RNAi flies (Figure S2F). Furthermore, *Lat>GABARAP-RNAi* males did not court target males in the dark (Figure S2E), indicating that male-male courtship defect is context-dependent and relies on the light conditions. Our results demonstrated that GABARAP function is required in Lat visual feedback neurons to suppress male courtship toward other males. Since *Lat>GABARAP-RNAi* males did not have gross anatomical abnormalities and the male-male courtship defect was sensory context-dependent, we hypothesized that GABARAP is required in Lat neurons for visual processing during courtship rather than regulating the development of male visual circuits.

**Figure 2.**
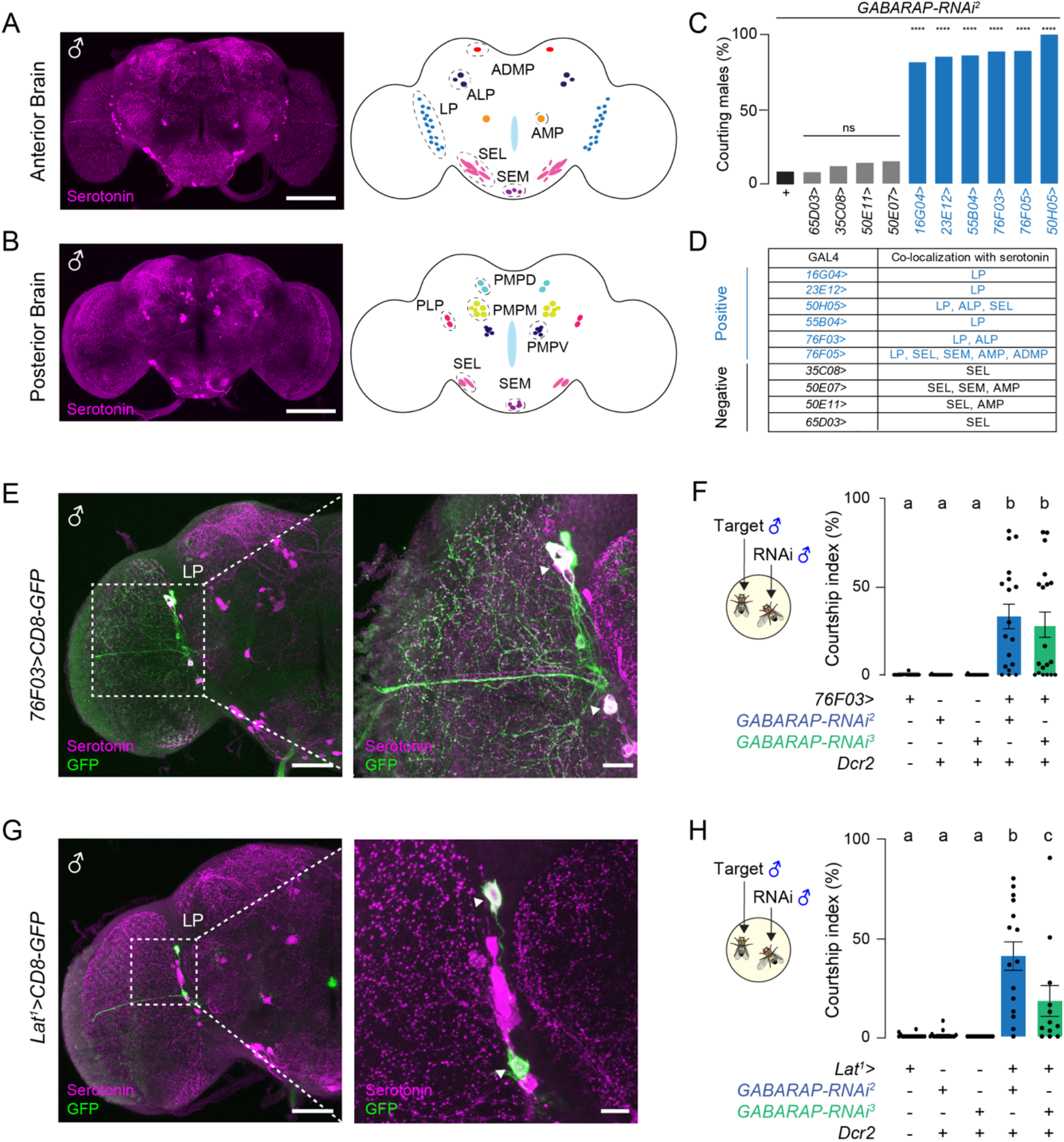
*GABARAP* knockdown in Lat neurons induces male courtship toward other males. (A-B) Serotonergic neurons in the anterior (A) and posterior (B) male brain (scale bars=50 μm). Illustration of serotonergic neurons in the central brain adapted from Pooryasin and Fiala, 2015. (C) Percentage of male-male courtship observed in various serotonergic GAL4s crossed to *GABARAP-RNAi*^*2*^. Positive GAL4s are colored in blue (Fisher’s exact test, ****, p<0.0001; n=23-58). (D) List of GAL4 strains tested and serotonergic neurons they label. (E) *76F03>CD8-GFP* male brains stained with anti-GFP (green) and anti-serotonin (magenta) (scale bars, left=50μm, right=10μm). White arrowheads indicate co-labeling. (F) Courtship quantification of *76F03*>*GABARAP-RNAi* and control males toward a male target (One-way ANOVA with Tukey’s test, p<0.001, mean ± SEM, n=16-18 flies). (G) *Lat*^*1*^*>CD8-GFP* male brains stained with anti-GFP (green) and anti-serotonin (magenta) (scale bars, left=50μm, right=10μm). White arrowheads indicate co-labeling. (H) Courtship quantification of *Lat*^*1*^>*GABARAP-RNAi* and control males toward a male target (One-way ANOVA with Tukey’s test, p<0.01, mean ± SEM, n=12-18 flies).

### The human ortholog of *Drosophila* GABARAP rescues the male-male courtship defect

In flies, GABARAP (Atg8a) functions have been studied mainly in the context of autophagy (Bali and Shravage, 2017; Jipa et al., 2021; Ratliff et al., 2015; Simonsen et al., 2008). However, mammalian GABARAPs are known to have autophagy-independent functions (Schaaf et al., 2016). To test whether GABARAP (Atg8a) function in Lat neurons is required for cellular autophagy, we checked for the accumulation of an autophagy disruption marker, Ref(2)P (Nezis et al., 2008), in *Lat>GABARAP-RNAi* males. Surprisingly, we did not detect an accumulation of Ref(2)P-positive protein aggregates in Lat neurons in RNAi males or controls (Figure S3A), indicating that GABARAP (Atg8a) in Lat neurons is not required for autophagy and GABARAP (Atg8a) might have autophagy-independent functions in these neurons. Mammalian GABARAPs have been shown to regulate the membrane trafficking and the synaptic localization of the GABA_A_ receptors (Kanematsu et al., 2007; Leil et al., 2004; Ye et al., 2021). In fact, before our study, no one has previously reported the strong sequence similarity between the *Drosophila* GABARAP (Dro-GABARAP) and the human GABARAP (hu-GABARAP) (Figures 3A and 3B). Here, we showed that Dro-GABARAP isoforms and the hu-GABARAP share remarkable sequence identities ranging from 68% to 88% at the amino acid level (Figure 3B). On the other hand, the amino acid identity between Dro-GABARAP and another human Atg8 protein, human LC3B (hu-LC3B), ranges from 25% to 29% (Figure 3B). These data indicate that Dro-GABARAP is more closely related to hu-GABARAP than hu-LC3B (Figure 3C). To test whether this strong sequence similarity between Dro-GABARAP and hu-GABARAP also reflects functional homology, we overexpressed hu-GABARAP or hu-LC3B in Lat neurons in *Lat>GABARAP-RNAi* males. As predicted by our protein sequence analysis, expression of hu-GABARAP but not hu-LC3B rescued Dro-GABARAP function in Lat neurons and suppressed male-male courtship (Figure 3D). Together, our results showed that Dro-GABARAP has autophagy-independent functions in Lat neurons, and hu-GABARAP can rescue this function, suggesting that Dro-GABARAP is a functional homolog of hu-GABARAP.

**Figure 3.**
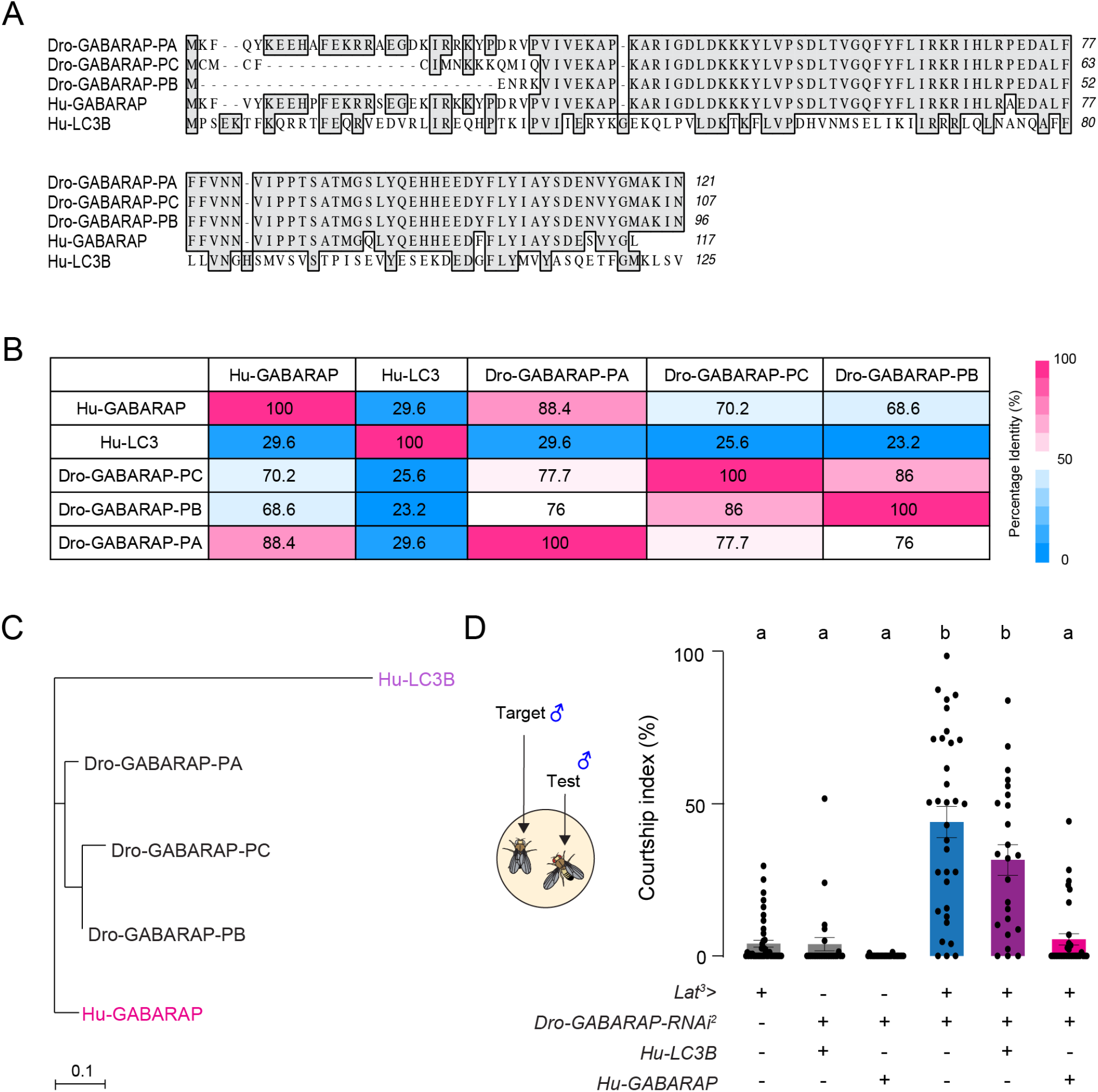
Human GABARAP can rescue *Drosophila* GABARAP function in Lat neurons. (A) Sequence alignment of *Drosophila melanogaster* GABARAP isoforms (GABARAP-PA/PB/PC) and their human orthologs, LC3B, GABARAP. Amino acid sequences were obtained from UniProtKB (www.uniprot.org) and aligned using T-Coffee multiple sequence alignment. Identical residues are highlighted in gray. (B) Table showing the percentage identity scores between *Drosophila* GABARAP isoforms and the human orthologs, LC3B and GABARAP. (C) Phylogenetic tree of *Drosophila* GABARAP and human orthologs, LC3B and GABARAP. The tree was constructed using the neighbor-joining method (MacVector). (D) Courtship quantification of males expressing *Lat*^*3*^*>GABARAP-RNAi*^*2*^ with and without human orthologs, *Hu-LC3B* or *Hu-GABARAP* toward a male target (One-way ANOVA with Tukey’s test, mean ± SEM, p<0.0001, n=24-44 flies).

### Hyperactivation of Lat neurons induces male courtship toward other males

In mammals, GABARAPs regulate neuronal activity by modulating the trafficking of GABA_A_ receptors to the synaptic membrane (Kanematsu et al., 2007; Leil et al., 2004; Ye et al., 2021). Because we found that Dro-GABARAP is a functional homolog of hu-GABARAP (Figure 3D), we hypothesize that it might regulate Lat activity by altering GABA_A_ receptor function (Figure 4A). In *Drosophila melanogaster*, three genes encode GABA_A_ receptor subunits: *resistance to dieldrin (Rdl), GABA and glycine-like receptor of Drosophila (Grd)*, and *ligand-gated chloride channel homolog 3* (*Lcch3*) (Buckingham et al., 1994; Gisselmann et al., 2004). Rdl is the best-characterized GABA_A_ receptor subunit in flies (Buckingham, 2005; Liu et al., 2007). To investigate whether GABA_A_ receptor function in Lat neurons is required to regulate male courtship, we knocked down *Rdl* in Lat neurons. *Lat>Rdl-RNAi* males courted other males at similar levels to *Lat>GABARAP-RNAi* males (Figure 4B), demonstrating that GABA_A_ receptor function indeed is required in Lat neurons to suppress male-male courtship.

**Figure 4.**
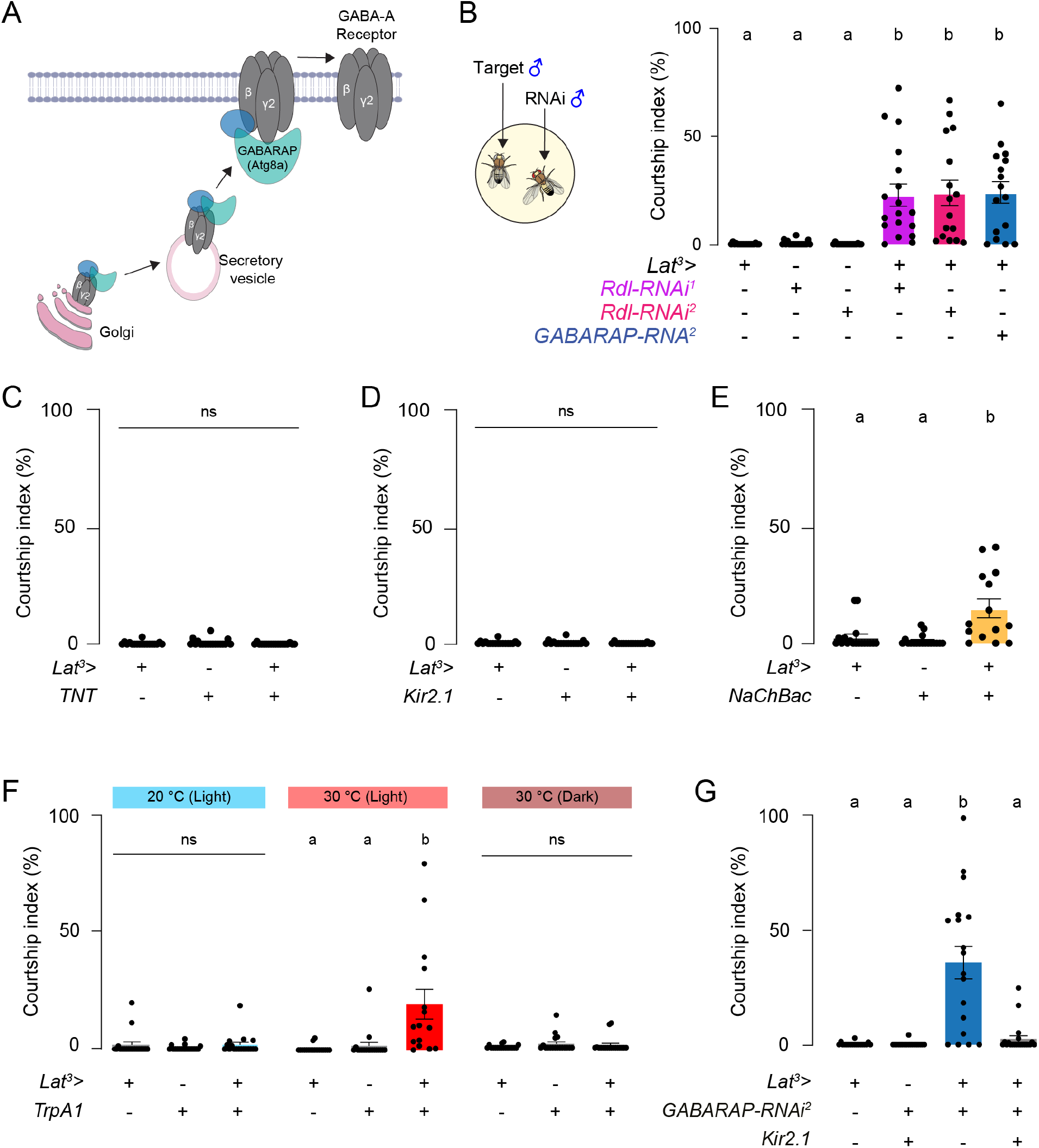
Hyperactivation of Lat neurons induces male courtship toward other males. (A) Model for GABA_A_ receptor membrane trafficking by GABARAP. Adapted from Kanematsu et al., 2007. (B) Courtship quantification of *Lat*^*3*^*>Rdl-RNAi* and control males toward a male target (One-way ANOVA with Tukey’s test, ns, mean ± SEM, n=16-18 flies). (C-D) Courtship quantification of *Lat*^*3*^*>TNT* (C), *Lat*^*3*^*>Kir2*.*1* (D), and control males toward a male target (One-way ANOVA with Tukey’s test, ns, mean ± SEM, n=18 flies). (E) Courtship quantification of *Lat*^*3*^*>NaChBac* and control males toward a male target (One-way ANOVA with Tukey’s test, p<0.001, mean ± SEM, n=14-18 flies). (F) Courtship quantification of *Lat*^*3*^*>TrpA1* and control males at 20°C and 30°C and in dark and light conditions toward a male target (One-way ANOVA with Tukey’s test, p<0.01, mean ± SEM, n=15-18 flies). (G) Courtship quantification of males expressing *Lat*^*3*^*>GABARAP-RNAi*^*2*^ with or without Kir2.1 toward a male target (One-way ANOVA with Tukey’s test, p<0.001, mean ± SEM, n=18-21 flies).

How do GABARAP and Rdl regulate Lat function? We speculated that GABARAP or GABA_A_ receptor function in Lat neurons is required to maintain Lat activity at a precise level to optimize male courtship. To test this, we genetically inhibited or activated Lat neurons. Silencing the activity of Lat neurons by expressing the tetanus toxin light chain (TNT) (Sweeney et al., 1995) (*Lat>TNT*, Figure 4C) or the Kir2.1 potassium channel (Baines et al., 2001; Johns et al., 1999) (*Lat>Kir2*.*1*, Figure 4D) did not induce male-male courtship. In contrast, chronic (Figure 4E) or temporal (Figure 4F) activation of Lat neurons by overexpressing the bacterial voltage-gated sodium channel, *NaChBac* (Ren et al., 2001), or transient receptor potential cation channel A1, *TrpA1* (Hamada et al., 2008) respectively, triggered male courtship toward other males. Similar to *Lat>GABARAP-RNAi* males, *Lat>TrpA1* males also did not court other males in the dark, once again demonstrating that the male-male courtship defect that originates in Lat neurons is context-dependent and is only induced under light conditions (Figure 4F, Video S4). We also tested *Lat>TNT* and *Lat>TrpA1* males in *Drosophila* activity monitors (DAM) typically used to measure circadian rhythms or sleep to investigate whether Lat neurons regulate these light-dependent behaviors. Manipulating Lat activity did not alter the daily phasic locomotor activity or sleep amount of male flies (Figures S4A-S4D). Finally, we activated other visual feedback neurons, Lawf1 and Lawf2, to test whether any visual perturbation can induce male-male courtship. Activation of Lawf1 or Lawf2 did not induce male-male courtship (Figures S4E-S4G), proving that courtship defect seen in *Lat>TrpA1* males is not caused by a general problem in males’ vision, and Lat neurons have a specific function in regulating male courtship.

Our data demonstrated that normal Lat activity is required to suppress male courtship toward other males under light conditions; when Lat neurons are hyperactivated, male flies abnormally court other males. Based on our results, we hypothesized that *GABARAP* knockdown in Lat neurons hyperactivates these neurons triggering male courtship toward other males. If this is true, silencing the activity of Lat neurons in *Lat>GABARAP-RNAi* males should suppress male courtship toward other males. To test this, we co-expressed *GABARAP-RNAi* and *Kir2*.*1* in Lat neurons and quantified male-male courtship. *Lat>GABARAP-RNAi* males courted other males but not when their activity was inhibited with Kir2.1 (Figure 4G). Together, our results suggest that Dro-GABARAP suppresses male-male courtship by preventing the hyperactivation of Lat neurons, potentially by altering GABA_A_ receptor signaling.

### Lat neurons are required for the vigorous following of females during courtship

We have shown that hyperactivation of Lat neurons triggers male courtship toward other males. However, this is an unusual behavior for the male flies because *Drosophila* males do not court other males in normal conditions. What is the normal function of Lat neurons in the male brain? Do Lat neurons regulate male courtship toward females? Since Lat neurons are visual feedback neurons, we speculated that they might modulate male visual processing during courtship. Visual input is critical for the ability of males to direct their locomotion and courtship actions toward females. Recently, a group of visual projection neurons, LC10, were shown to regulate the ability of males to track females during courtship (Hindmarsh Sten et al., 2021; Ribeiro et al., 2018). To investigate whether Lat neurons contribute to female tracking, we silenced their activity during male-female courtship (Figure 5A). Blocking synaptic release in either Lat or LC10 neurons significantly decreased male copulation success (Figure 5B). To further identify which courtship steps Lat neurons are required for, we tracked male and female behaviors at high resolution during courtship using a deep-learning-based framework for multi-animal pose tracking, SLEAP (Figure 5A and Video S5) (Pereira et al., 2019; Pereira et al., 2022). To train the SLEAP neural network, we labeled specific body parts (eyes, head, thorax, wing tips, and abdominal tip) of freely interacting pairs of male and female flies. After several training rounds, we confirmed that the SLEAP neural network effectively tracked male and female actions during courtship. Using the tracking data, we automatically classified specific male behaviors such as orientation, wing extension, and following (Figures 5C and 5D). The duration and the sequence of each male courtship action were then visualized using ethograms in *Lat>TNT, LC10>TNT*, and control flies allowing us to determine the specific courtship defects as a consequence of Lat or LC10 neural inhibition (Figure 5D). In control flies, the distributions of the male’s head angle and the distance between the male and female flies were biased toward small values. As previously shown by Ribeiro et al., 2018, inhibiting LC10 neurons increased the head angle and the distance between male and female flies during courtship (Figures 5E-5G). The ability of *Lat>TNT* males to orient toward females was not impaired; however, their distance from the female fly was slightly elevated during courtship (Figures 5E-5G). We further calculated the wing extension and the following index for *Lat>TNT, LC10>TNT*, and control flies. *LC10>TNT* males showed a decrease in both metrics toward females (Figures 5H and 5I), while *Lat>TNT* males were only impaired in following behavior (Figure 5I). Our data suggest that the activity of Lat neurons, similar to LC10 neurons, is required for the ability of males to track females during courtship. When these neurons are inhibited, males cannot vigorously follow females, and their copulation success is reduced.

**Figure 5.**
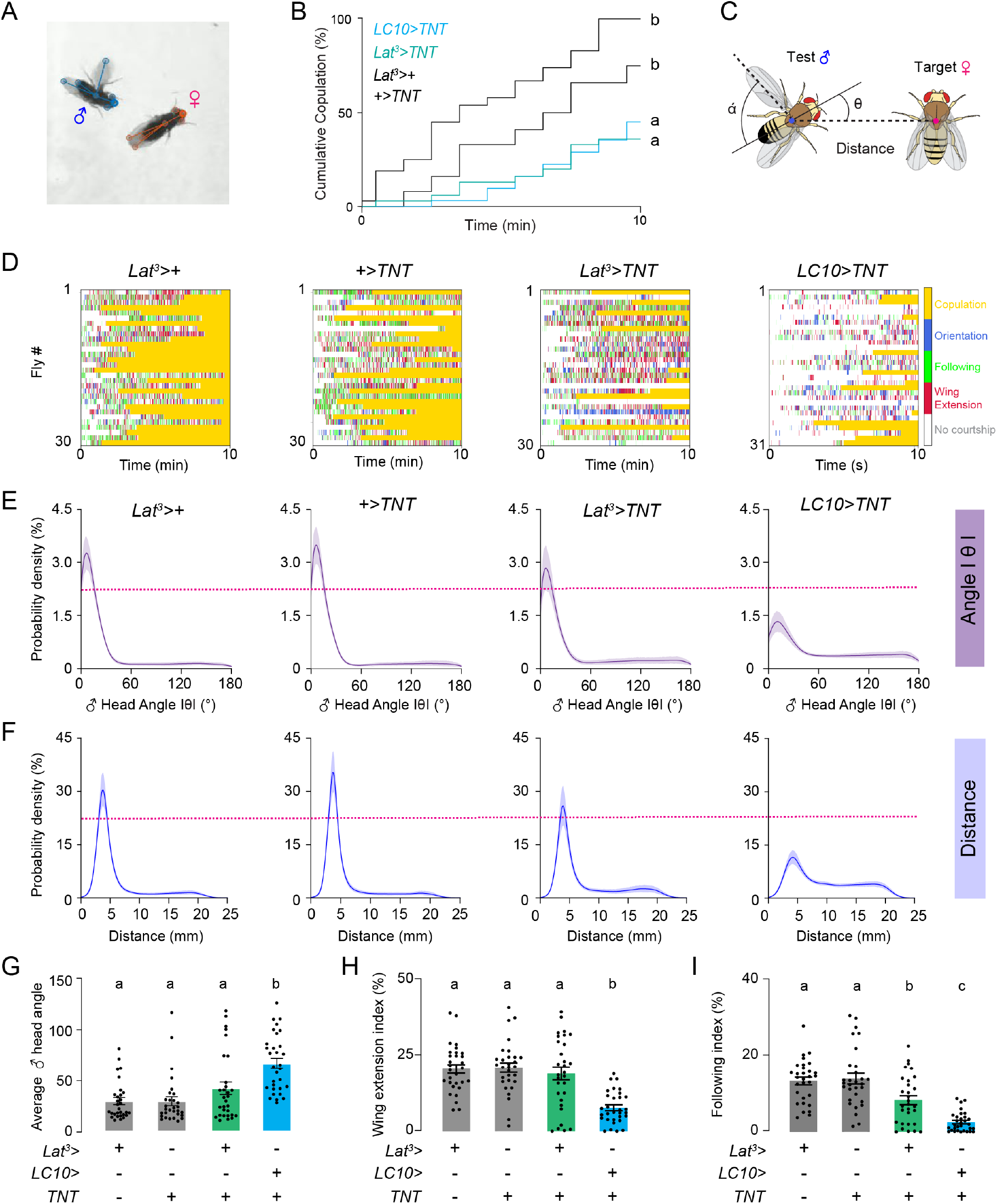
Inhibition of Lat neurons impairs the ability of males to follow females during courtship. (A) Snapshot of male (blue) and female (orange) fly tracking in SLEAP. (B) Cumulative percentage of males with indicated genotypes copulating in 10min (Kaplan-Meier, p <0.0001; mean ± SEM), (n=30-31 flies for all the analyses here and below). (C) Schematic showing head angle (θ), wing angle (α), and the distance between the male and the female. (D) Male courtship ethograms for indicated genotypes are generated using SLEAP tracking data. (E-F) Probability density functions of the absolute male head angle δ (E) and distance between male and female flies (F) of indicated genotypes during single-pair courtship. (G-I) Average head angle IθI (G), wing extension index (H), and following index (I) plotted for males of indicated genotypes (One-way ANOVA with Tukey’s test, p<0.05, mean ± SEM).

### Lat neurons impact male behavior via neural circuits that drive visually-guided courtship pursuits

Our genetic manipulations revealed that GABARAP function in Lat neurons is required to optimize their activity and stimulate or suppress male courtship. To further understand how these neurons regulate male behavior, we investigated their anatomical organization and functional connections in detail. Lat neurons are present in the brains of both sexes (Kolodziejczyk, 2011; Kolodziejczyk and Nässel, 2011b). Lat cell bodies are located between the optic lobe and the central brain (Figures 6A and 6B). These neurons are proposed to provide feedback from the central brain to the distal surfaces of the lamina (Kolodziejczyk and Nässel, 2011b). Consistent with previous studies, we found that Lat axon terminals labeled by *Lat>Syt-GFP* (Zhang et al., 2002) are located in the lamina (Figure 6C), and Lat dendritic arbors labeled by *Lat>DenMark* (Nicolai et al., 2010) are located in the posterolateral protocerebrum (Figure 6D). To complement our light microscopy analysis and to further investigate Lat pre- and post-synaptic sites and their connectivity, we used FlyWire, an open-source connectomic resource for the female *Drosophila* brain (Dorkenwald et al., 2022). We identified and proofread 12 putative Lat neurons within the electron microscopic (EM) brain volume (Zheng et al., 2018) and reconstructed them using a standard fly brain template (Figure 6E, Table S1, and Video S6). Our EM tracing results identified three types of Lat neurons: Lat-Type1 neurons innervate both dorsal and ventral segments of the lamina (Figures 6F and S5A). In contrast, Lat-Type2 and Lat-Type3 neurons arborize in the dorsal or ventral lamina, respectively (Figures 6F, S5B, and S5C). We also used the EM data to determine the number of synaptic connections between putative Lat neuron subtypes via an automated synapse detection algorithm (Buhmann et al., 2021). Using the connectivity data, we built a neural connectivity matrix and showed that there is little or no connectivity between 12 putative Lat neurons, with one exception; Lat3 neurons are presynaptic to Lat6 neurons (8 synapses) (Figure 6G). We next investigated the neurotransmitters used by Lat neurons using specific antibodies for dopaminergic, octopaminergic/tyraminergic, serotonergic, and GABAergic neurons. We found that Lat neurons with cell bodies in the dorsal protocerebrum co-expressed markers for serotonin and octopamine/tyramine, whereas Lat neurons with cell bodies in the ventral protocerebrum co-expressed markers for dopamine and octopamine/tyramine (Figures S6A-S6C). We did not detect markers for cholinergic or GABAergic neurons in any of the Lat neurons identified (Figures S6D and S6E). Together these results demonstrate that Lat neurons are modulatory visual feedback neurons that receive input from the posterolateral protocerebrum in the central brain and relay information to the distal regions of the lamina. They can be classified into different subtypes based on their axonal arborizations in the lamina and the neurotransmitters they express.

**Figure 6.**
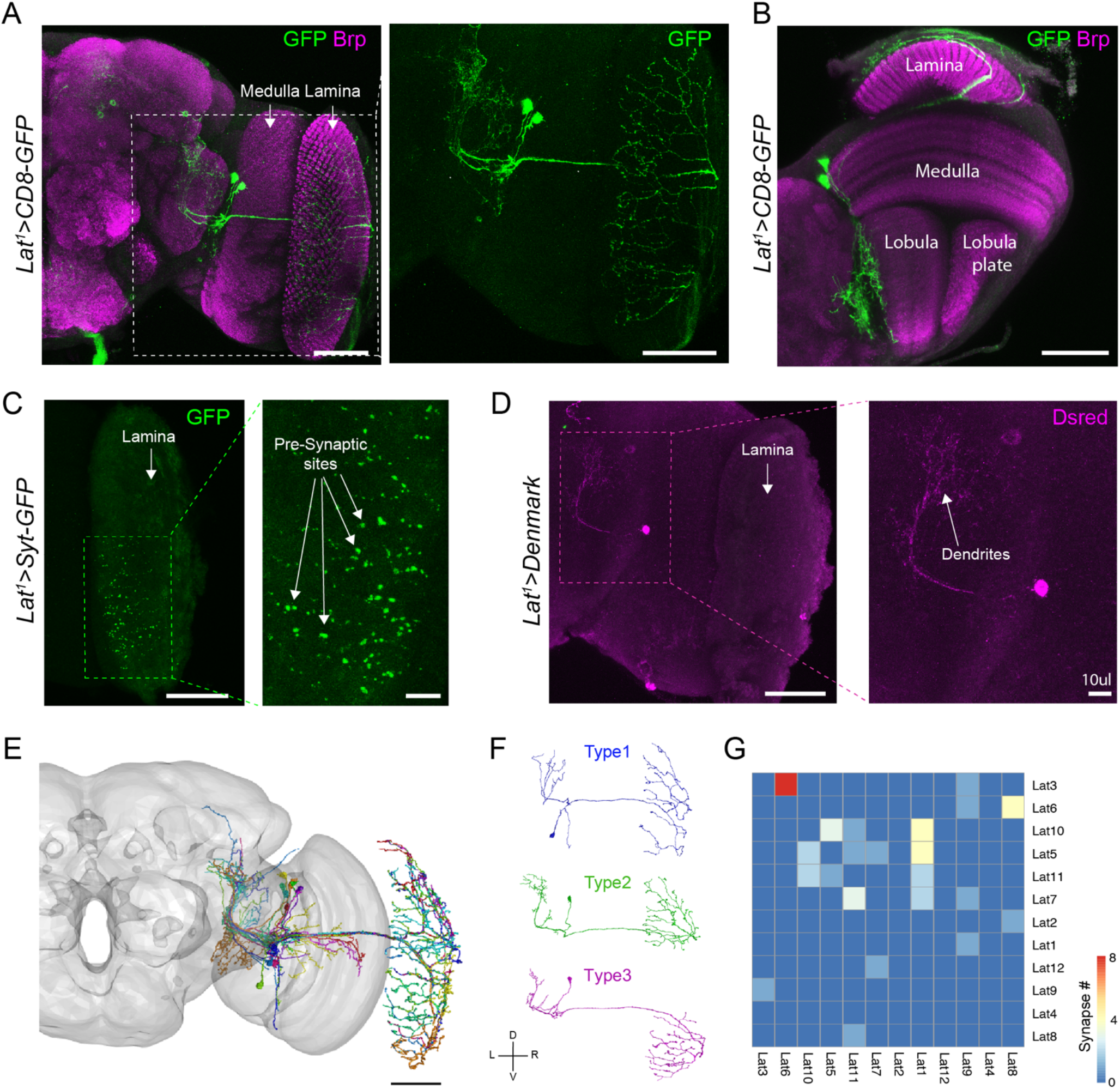
Lat neurons have different subtypes. (A) *Lat*^*1*^*>CD8-GFP* male brain stained with anti-GFP (green) and anti-Brp (magenta). Labels indicate different regions of the optic lobe. The right panel shows magnified Lat neurons (scale bars=50μm). (B) *Lat*^*1*^*>CD8-GFP* male brain showing the lamina and neuropil layers of the optic lobe (scale bars=50μm). (C) *Lat*^*1*^*>Syt-GFP* male brain stained with anti-GFP (green) showing axonal terminals of Lat neurons in the lamina (scale bars left=50μm, right=10μm). (D) *Lat*^*1*^*>DenMark* male brain stained with anti-DsRed (magenta) showing dendrites of Lat neurons in the lateral protocerebrum (scale bars left=50μm, right=10μm). (E) Reconstruction of putative Lat neurons (n=12) in the right hemisphere using the Flywire segmentation of an electron microscopic (EM) volume of an entire female fly brain (gray) (scale bars left=50μm). (F) We identified three classes of Lat neurons based on axonal projection patterns in the lamina. Example EM reconstructions of Lat-Type1, Lat-Type2, and Lat-Type3 neurons are shown. (G) Heatmap representing the synaptic connectivity matrix across putative Lat neurons.

Next, to identify the neural circuit mechanisms by which Lat neurons regulate male behavior, we investigated their functional connectivity to neurons that are previously shown to mediate visually-guided male courtship, LC10 (Hindmarsh Sten et al., 2021; Ribeiro et al., 2018), and Fru-P1 neurons (Kimura et al., 2008; Kohatsu et al., 2011; Von Philipsborn et al., 2011). LC10 neurons have four distinct subtypes, LC10a-d, classified by their arborization patterns in the lobula (Wu et al., 2016). LC10a activity is particularly critical for visually-guided male courtship pursuits (Hindmarsh Sten et al., 2021; Ribeiro et al., 2018). To examine the functional connections between Lat and LC10a neurons, we used *in vivo* optogenetic stimulation coupled with two-photon calcium imaging (Figure 7A). We first stimulated Lat neurons while imaging the activity of LC10a neurons using a genetically encoded calcium indicator GCaMP6s (Chen et al., 2013). Lat activation caused a robust increase in GCaMP6s fluorescence in LC10a neurons (Figures 7B-7D) but not in other visual projection neurons tested (Figures S7A-S7D). Next, we optogenetically stimulated LC10a-d neurons while recording the activity of Lat neurons. Optogenetic activation of LC10a-d neurons elicited calcium responses in Lat neurons, but these responses were not significantly different from controls (Figures S7E-S7H). Our results demonstrated that Lat neurons provide functional input to LC10a neurons, but LC10a-d neurons do not provide functional input to Lat neurons. How does the Lat>LC10a pathway regulate male courtship? LC10a activity is strongly enhanced when males are in a courtship state or when male-specific Fru-P1 neurons are activated (Hindmarsh Sten et al., 2021; Ribeiro et al., 2018). We hypothesized that the Lat>LC10a pathway might regulate male courtship via their interactions with Fru-P1 neurons. Indeed, optogenetic stimulation of LC10a neurons was sufficient to induce calcium responses in Fru-P1 neurons, demonstrating that the Lat-LC10a pathway is functionally connected to Fru-P1 neurons (Figures 7E-7H).

**Figure 7.**
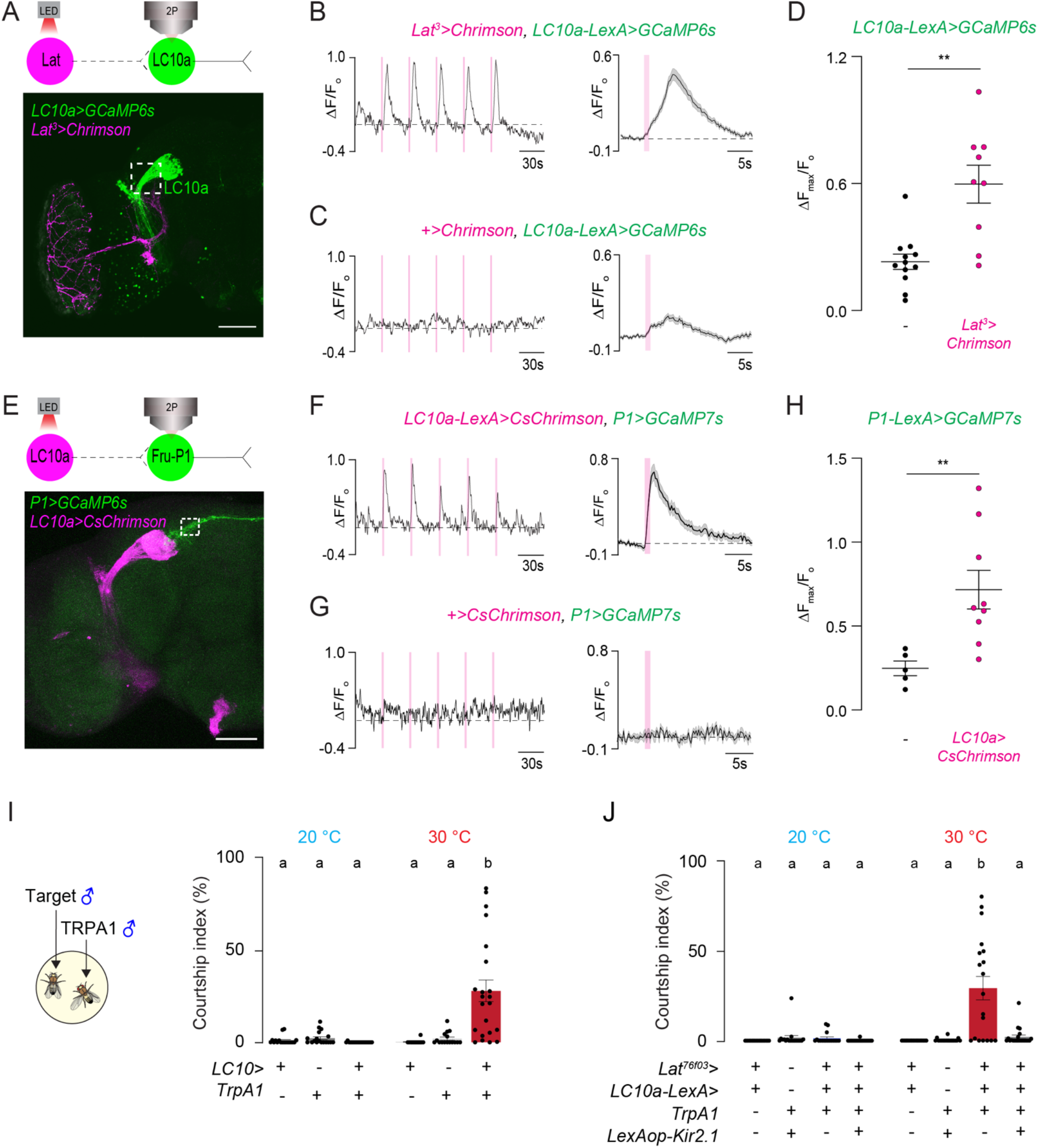
Lat neurons are functionally connected to visual circuits that regulate courtship pursuits. (A) Schematic of the *in viv*o optogenetic stimulation/two-photon imaging setup (top). *Lat*^*3*^*>Chrimson* (magenta), *LC10a-LexA>GCaMP6s* (green) example brain (scale bar= 50 μm, white box=ROI). (B) Individual (left) and averaged (right) normalized responses of LC10a neurons upon optogenetic stimulation of Lat neurons (mean ± SEM; n=9 flies, 1s pulsed (20Hz) LED stimulation). (C) Individual (left) and averaged (right) normalized responses of LC10a neurons in control flies upon optogenetic stimulation (mean ± SEM; n=12 flies, 1s pulsed (20Hz) LED stimulation). (D) Averaged normalized peak responses of LC10a neurons in indicated genotypes upon optogenetic stimulation of Lat neurons (Unpaired t-test with Welch’s correction; p<0.01; mean ± SEM, n=9-12 flies). (E) Schematic of the *in vivo* optogenetic stimulation/two-photon imaging setup (top). *LC10a-LexA>CsChrimson* (magenta), *P1>GCaMP7s* (green) example brain (scale bar=50 μm, white box=ROI) (F) Individual (left) and averaged (right) normalized responses of P1 neurons upon optogenetic stimulation of LC10a neurons (mean ± SEM; n=9 flies, 1s pulsed (20Hz) LED stimulation). (G) Individual (left) and averaged (right) normalized responses of P1 neurons in control flies upon optogenetic stimulation (mean ± SEM; n=5 flies, 1s pulsed (20Hz) LED stimulation). (H) Averaged normalized peak responses of P1 neurons in indicated genotypes upon optogenetic stimulation of LC10a neurons (Unpaired t-test with Welch’s correction; p<0.01; mean ± SEM, n=5-9 flies). (I) Courtship quantification of *LC10>TrpA1* and control males toward a male target at 20°C (n=18 flies) and 30°C (n=16-22 flies) (One-way ANOVA with Tukey’s test, p<0.001, mean ± SEM). (J) Courtship quantification in indicated genotypes at 20°C (n=17-20 flies) and 30°C (n=18-24 flies). Silencing LC10a neurons during thermogenetic activation of Lat neurons represses courtship toward a male target. (One-way ANOVA with Tukey’s test, p<0.0001, mean ± SEM).

Our functional imaging experiments suggest that Lat neurons regulate male courtship behavior, specifically the ability of males to track females via their interactions with the LC10a-Fru-P1 neurons. If this is true, activation of LC10a-d neurons should also trigger male-male courtship, and Lat-induced male-male courtship should rely upon LC10a activation. As our model predicted, activation of LC10a-d neurons induced male-male courtship (Figure 7I), and silencing of the activity of LC10a neurons during Lat thermogenetic activation suppressed male-male courtship (Figure 7J). Taken together, our results clearly demonstrated that Lat neurons regulate male courtship via the LC10a-Fru-P1 pathway.

## Discussion

*Drosophila melanogaster* males use multi-modal sensory information to identify and mate with suitable females. Although neural circuits that process olfactory, gustatory, and auditory cues during courtship have been extensively studied (Baker et al., 2022; Clowney et al., 2015; Deutsch et al., 2019; Dickson, 2008; Kallman et al., 2015; Kohl et al., 2013; Kurtovic et al., 2007; Thistle et al., 2012; Von Philipsborn et al., 2011), neural circuits that process visual information are just beginning to be characterized. In this study, we revealed that a *Drosophila* ortholog of human GABA_A_-receptor-associated protein (GABARAP) is required in a small population of visual feedback neurons, Lat, to optimize male courtship behavior. Knocking down *GABARAP* or *GABA*_*A*_ *receptors* in Lat neurons induces male-male courtship while inhibiting their activity impairs male-female courtship. Lat neurons are functionally connected to LC10a visual projection neurons which output to Fru-P1 neurons to modulate male courtship. We propose a model in which GABARAP/GABA_A_ receptor signaling maintains Lat neuron activity in a precise regime: Activation above this regime hyperactivates the LC10a-Fru-P1 circuit, reducing male fly’s ability to discriminate males from females and leading to male-male courtship. Inhibiting Lat activity, on the other hand, below this regime impairs male-female courtship as if the visual system is now not sensitive enough to track the female fly.

### GABARAP regulates Lat activity via GABA_A_ receptor signaling

Previous studies have focused mainly on studying the functions of *Drosophila* GABARAP (Atg8a) in the cellular degradation pathway autophagy (Bali and Shravage, 2017; Jipa et al., 2021; Ratliff et al., 2015; Simonsen et al., 2008). However, these studies have not realized that GABARAP (Atg8a) shares up to 88% sequence identity with the mammalian GABARAPs, which regulate the surface expression and membrane trafficking of GABA_A_ receptors (Leil et al., 2004; Ye et al., 2021). In addition to GABARAP (Atg8a), other Atg proteins are also shown to play essential roles in regulating neuronal physiology and behavior (Hui et al., 2019; Lieberman et al., 2020; Nikoletopoulou and Tavernarakis, 2018; Schaaf et al., 2016; Yang et al., 2022). For example, in the mouse striatum, conditional knockouts of *Atg7* in two different types of GABAergic neurons; (dSPN and iSPN) lead to defects in motor learning by either impairing the dendritic arborization or the resting membrane potential of these neurons (Lieberman et al., 2020). In the forebrain GABAergic neurons, *Atg7* knockouts disrupt social behaviors. The behavioral impairments seen in *Atg7* mutants are due to the reduction of surface GABA_A_ receptor levels and the hyperactivation of GABAergic neurons (Hui et al., 2019). These studies suggest that Atg proteins in the nervous system have autophagy-independent functions, and they regulate neuronal physiology and animal behavior by modulating GABA_A_ receptor signaling.

Our study found that Dro-GABARAP and GABA_A_ receptor functions are required in a group of visual feedback neurons, Lat, to suppress male courtship toward other males. Strikingly, the function of Dro-GABARAP in Lat neurons can be rescued by the expression of its human ortholog (Figure 3). Our findings support the idea that similar to the mammalian systems, Dro-GABARAP in flies might regulate neural activity by modulating GABA_A_ receptor localization at the synaptic terminals. The results of our neuronal activation experiments further support this model (Figures 4E and 4F). It is still possible that Dro-GABARAP function in Lat neurons is also required for maintaining the structural integrity of these neurons. Although we have not found evidence for autophagy disruption in Lat neurons upon *Dro-GABARAP* knockdown (Figure S3A), it might regulate other cellular pathways in Lat neurons that we have not characterized. Future experiments should investigate the consequences of Dro-GABARAP loss of function in Lat neurons by transcriptomic and proteomic analysis and determine which molecular pathways are disrupted. Another piece of evidence supporting our model that Dro-GABARAP regulates Lat neural activity rather than its structural organization is based on our anatomical characterization of Lat neurons. We did not observe any gross abnormalities in the dendritic or synaptic arborization patterns of Lat neurons upon RNAi-mediated knockdown of *Dro-GABARAP* (Figures S2F and S2G). In addition, *Dro-GABARAP* knockdown-induced male-male courtship was reversible and dependent on the environment (Figure S2E), suggesting Dro-GABARAP likely modulates the activity of Lat neurons rather than their wiring or cellular homeostasis.

### Lat neurons are visual feedback neurons that connect the central brain to the lamina

Lat neurons (also referred to as LBO5HT) were first described as serotonin-immunoreactive cells that send processes to the distal lamina in *Drosophila* and other insects (Kolodziejczyk and Nässel, 2011a, b; Nassel, 1988; Nässel et al., 1987; Pyza and Meinertzhagen, 1996). Since their first characterization in the 1980s, the functions of Lat neurons in visual processing and behavior have remained poorly understood. Recent studies aiming to characterize the roles of lamina-associated neurons in response to a diverse set of motion stimuli found no behavioral defects in flies where Lat neurons were activated or inhibited (Tuthill et al., 2013). Our study is the first to unveil a potential role for Lat neurons in regulating visual processing, specifically during *Drosophila* male courtship behavior.

In blowflies, *Calliphora erythrocephala*, Lat neurons do not possess canonical presynaptic sites in the lamina (Nässel and Elekes, 1984). In *Drosophila melanogaster*, axon terminals of Lat neurons do not express presynaptic markers such as Bruchpilot or synapsin (Kolodziejczyk, 2011). These earlier observations and our current EM analysis led to the idea that Lat neurons might release neurotransmitters in the lamina in a paracrine fashion. In our study, we found that Lat neurons express multiple neurotransmitters in addition to serotonin; dorsal Lat neurons are both serotonergic and octopaminergic, and ventral Lat neurons are both dopaminergic and octopaminergic (Figures S6A-S6C). Co-expression of different neurotransmitters in the *Drosophila* brain has been previously reported (Deng et al., 2019). We have not characterized when or how each neurotransmitter is released from Lat neurons or their particular functions. However, based on their anatomical organization and the neurotransmitter pairs they express, we speculate that dorsal and ventral Lat neurons might have distinct roles in regulating fly visual processing during courtship or other visually-guided behaviors.

Here we focused on the functions of Lat neurons in male flies. But Lat neurons exist in both sexes. The function of Lat neurons in females is currently unknown, and we did not observe any apparent defects in *Trh>GABARAP-RNAi* females in our experiments (Figure 1F). In male flies, we showed that Lat neurons are required for visually-guided tracking behavior during courtship. However, we do not know which sensory signals Lat neurons respond to. In fact, based on their anatomical organization, we propose that Lat neurons might not directly respond to visual features or motion for two reasons: 1. Most of the Lat dendrites are not located in the optic lobe but in the central brain. 2. Lat neurons are not connected to photoreceptors or other visually-responsive neurons in the lamina (Nässel and Elekes, 1984; Rivera-Alba et al., 2011). We speculate that Lat neurons respond to internally-generated signals and relay them from the central brain back to the lamina to adjust early visual processing during courtship.

### Lat neurons regulate male courtship by altering the activity of LC10a visual projection neurons

Our data demonstrate that Lat neurons regulate male courtship through their functional connections with the LC10a-Fru-P1 circuit (Figure 7). The functional connections between Lat and LC10a neurons are not direct since our EM analysis did not identify any synaptic connections between Lat and LC10a neurons in the female connectome. However, we could not assess the synaptic connectivity between Lat and LC10a neurons in the male brain because a male connectome does not exist yet. Still, based on the anatomical organization of these neurons (Figure 7A), we think the functional connectivity between Lat and LC10a neurons must be multisynaptic. Furthermore, our study demonstrated that optogenetic stimulation of LC10a neurons is sufficient to activate Fru-P1 neurons. Previous studies have shown that Fru-P1 neurons regulate LC10a responses during tracking based on the male’s sexual arousal state (Hindmarsh Sten et al., 2021). Our optogenetic activation experiments showed that LC10a neurons provide excitatory input to Fru-P1 neurons, but these neurons also do not share direct synaptic connections (Hindmarsh Sten et al., 2021; Ribeiro et al., 2018). We currently do not know how Lat neurons interact with LC10a and Fru-P1 neurons. We speculate that LC10a and Lat neurons might generate feed-forward excitation in the visual system to enhance male visual perception. Activation of Lat neurons by currently unknown factors stimulates LC10a neurons which then will activate Fru-P1; activation of Fru-P1 will further boost LC10a activity and allow the male fly to rigorously track the moving female target. Future studies should record the activity of these neurons simultaneously when a male fly is engaged in courtship pursuits to reveal the dynamic interactions between them.

### A conserved mechanism for regulating visual processing during mating behaviors

The data presented here provide strong evidence that GABARAP/GABA_A_ receptor signaling in the visual feedback neurons Lat contributes to the mating success of *Drosophila* males by modulating their visual perception. Based on the structural and functional similarities between the insect and vertebrate visual systems, we speculate that GABARAP and GABA_A_ receptors might have conserved functions in modulating the activity of visual circuits and social behaviors in vertebrates as well as in insects. In fact, efferent fibers entering the retina, retinopetal fibers, have been found in many vertebrate species (Ball et al., 1989; Brooke et al., 1965; Cajal, 1892; Halpern et al., 1976; Schütte, 1995; Uchiyama, 1989). Similar to Lat neurons in flies, in rodents, some of the retinopetal fibers are serotonergic and originate from the dorsal raphe nucleus (DRN) of the brainstem (Repérant et al., 2000; Villar et al., 1987). Stimulation of DRN efferents enhances visual responses in the retina (Lörincz et al., 2008), but the behavioral consequences of this visual enhancement remain poorly understood. A recent study showed that retinal output is modulated by arousal in mice (Schröder et al., 2020). Whether the arousal state modulates the visual gain through the activation of retinopetal fibers is unclear. Given that Lat neurons are visual feedback neurons that project to the posterior lamina right below the fly retina, we speculate that Lat neurons in flies and retinopetal fibers in vertebrates regulate early visual processing by providing feedback signals to the retina when animals are in different arousal states.

## Acknowledgments

We thank Mariana Wolfner, Inês Ribeiro, Madineh Sarvestani, Vanessa Ruta, Joseph Fetcho, Leslie Vosshall, and members of the Yapici Lab for their comments on the manuscript. We thank Kathi Eichler for her assistance with EM data analysis. We thank Vanessa Ruta and Tom Hindmarsh Sten for their advice on LC10a two-photon imaging. We thank Caleb Vogt for his assistance in using the SLEAP software. Y.M. is supported by Funai Overseas Scholarship. Research in N.Y.’s laboratory is supported by a Cornell University Nancy and Peter Meinig Family Investigator Program, a Pew Biomedical Scholar Award, the Alfred P. Sloan Foundation Award, and NIH R35 ESI-MIRA Grant (R35GM133698-01). We acknowledge Bloomington Drosophila Stock Center (NIH P40OD018537) and the Developmental Studies Hybridoma Bank (NICHD of the NIH, University of Iowa) for reagents. Imaging data were acquired through the Cornell University Biotechnology Resource Center, with NIH S10OD018516 funding for the shared Zeiss LSM880 confocal/multiphoton microscope. We acknowledge the Princeton FlyWire team and members of the Murthy and Seung labs for the development and maintenance of FlyWire (supported by BRAIN Initiative grant MH117815 to Murthy and Seung). Ongoing development of the natverse, including the fafbseg package, is supported by the NIH BRAIN Initiative (grant 1RF1MH120679-01) and the Medical Research Council (MC_U105188491).

## Author contributions

Y.M. and N.Y. conceived the project and designed all the experiments. Y.M. carried out and analyzed all the experiments. X.C. helped with two-photon imaging analysis and proofreading of Lat neurons in the Flywire. L.X. helped with experiments in Figure S1. H.K. built the chamber in Figure 6 and prepared the supplementary videos. T.J. wrote the code for plotting two-photon imaging data. Y.M. and N.Y. interpreted the results and wrote the paper with feedback from all authors.

## Supplementary figures and legends

**Figure S1.**
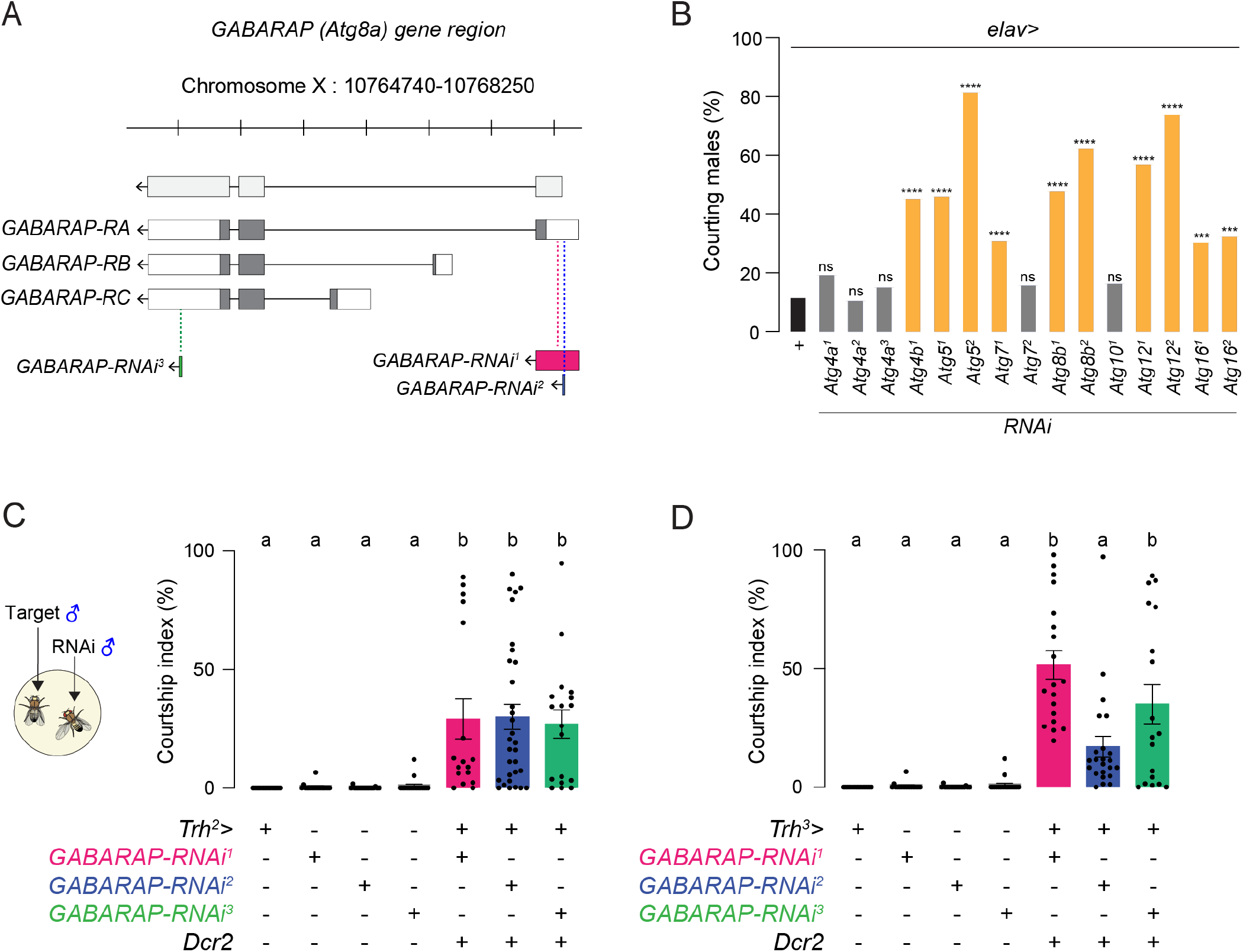
*GABARAP* knockdown in serotonergic neurons induces male-male courtship, related to Figure 1. (A) Schematic showing the *GABARAP* (*Atg8a*) gene locus in the *Drosophila* genome and the location of various RNAi hairpins targeting different regions of the *GABARAP* gene. (B) Percentage of males courting a male target in various *GABARAP-RNAi* lines crossed to *elav-GAL4*. Positive RNAi lines are colored in orange (Fisher’s exact test, ***p<0.001, ****p<0.0001; n=83-153). (C) Courtship quantification of *Trh*^*2*^*>GABARAP-RNAi* and control males toward male targets (One-way ANOVA with Tukey’s test, p<0.01, mean ± SEM, n=17-31 flies). (D) Courtship quantification of *Trh*^*3*^*>GABARAP-RNAi* and control males toward male targets (One-way ANOVA with Tukey’s test, p<0.05, mean ± SEM, n=18-23 flies). The same control data for RNAi lines in Figure 1E are used in (C) and (D) because these courtship assays were performed on the same days in parallel but plotted separately.

**Figure S2.**
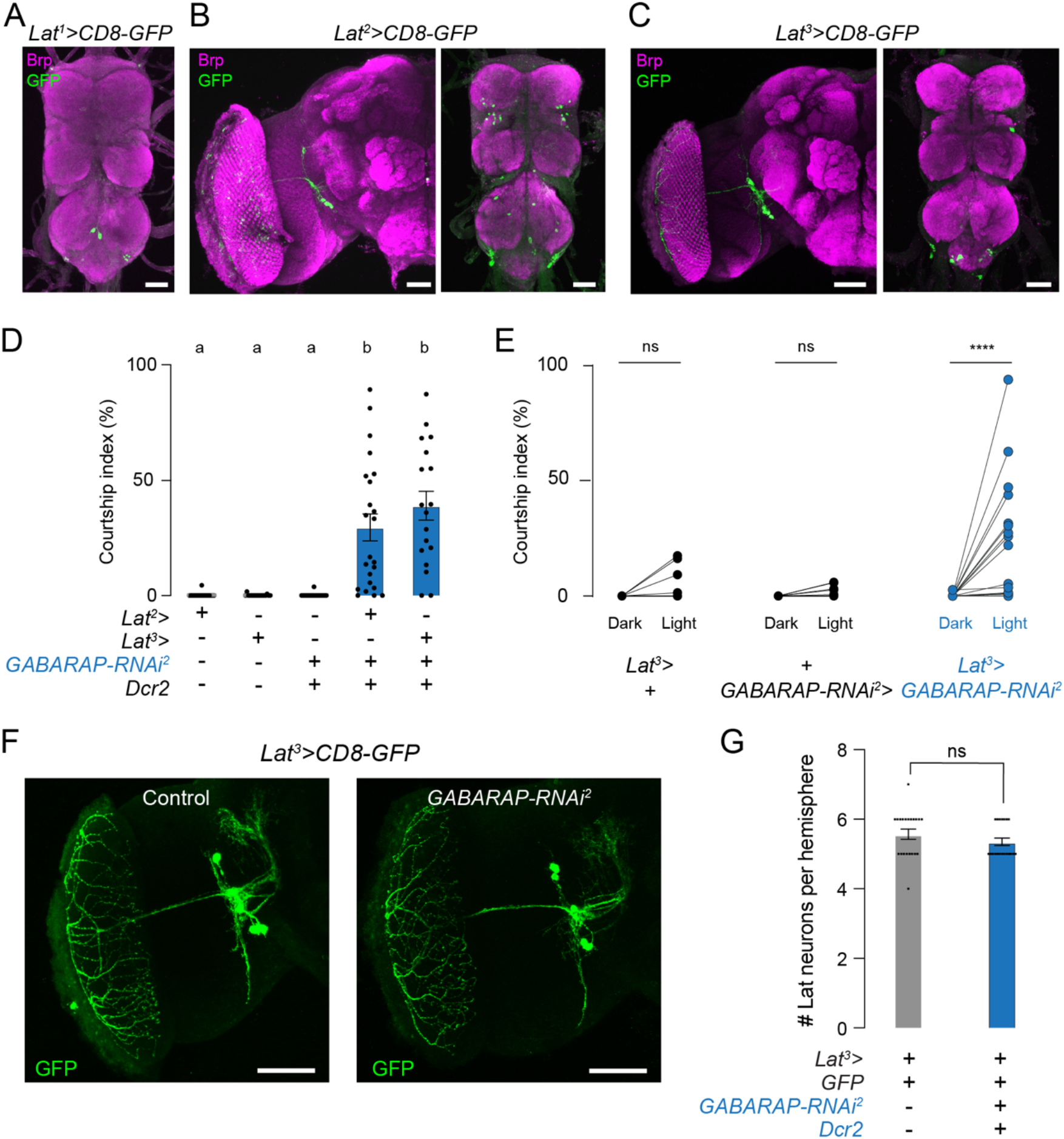
*GABARAP* knockdown in Lat neurons induces male-male courtship only in light conditions, and it does not impact the number of Lat neurons in the brain, related to Figure 2. *(A) Lat*^*1*^*>CD8-GFP* male VNC stained with anti-GFP (green) and anti-Brp (magenta) (scale bars=10μm). *(B) Lat*^*2*^*>CD8-GFP* male brain and VNC stained with anti-GFP (green) and anti-Brp (magenta) (scale bars; brain=50μm, VNC=10μm). *(C) Lat*^*3*^*>CD8-GFP* male brain and VNC stained with anti-GFP (green) and anti-Brp (magenta) (scale bars; brain=50μm, VNC=10μm). (D) Courtship quantification of *Lat*^*2*^*>GABARAP-RNAi*^*2*^, *Lat*^*3*^*>GABARAP-RNAi*^*2*,^ and control males toward a male target (One-way ANOVA with Tukey’s test, p<0.0001, mean ± SEM, n=18-23 flies). (E) Courtship quantification of *Lat*^*3*^*>GABARAP-RNAi*^*2*^, and control males toward a male target in the light and dark conditions (Wilcoxon matched-pairs signed rank test, p<0.0001, n= 17-18 flies). *(F) Lat*^*3*^*>CD8-GFP* brains with and without *GABARAP-RNAi* expression are stained with anti-GFP (green) to determine potential anatomical defects in Lat neurons (scale bars=50μm). (G) Quantification of the number of Lat cell bodies in *Lat*^*3*^*>CD8-GFP* flies with and without *GABARAP-RNAi* expression (Unpaired t-test with Welch’s correction; ns, mean ± SEM, n=10-11 flies).

**Figure S3.**
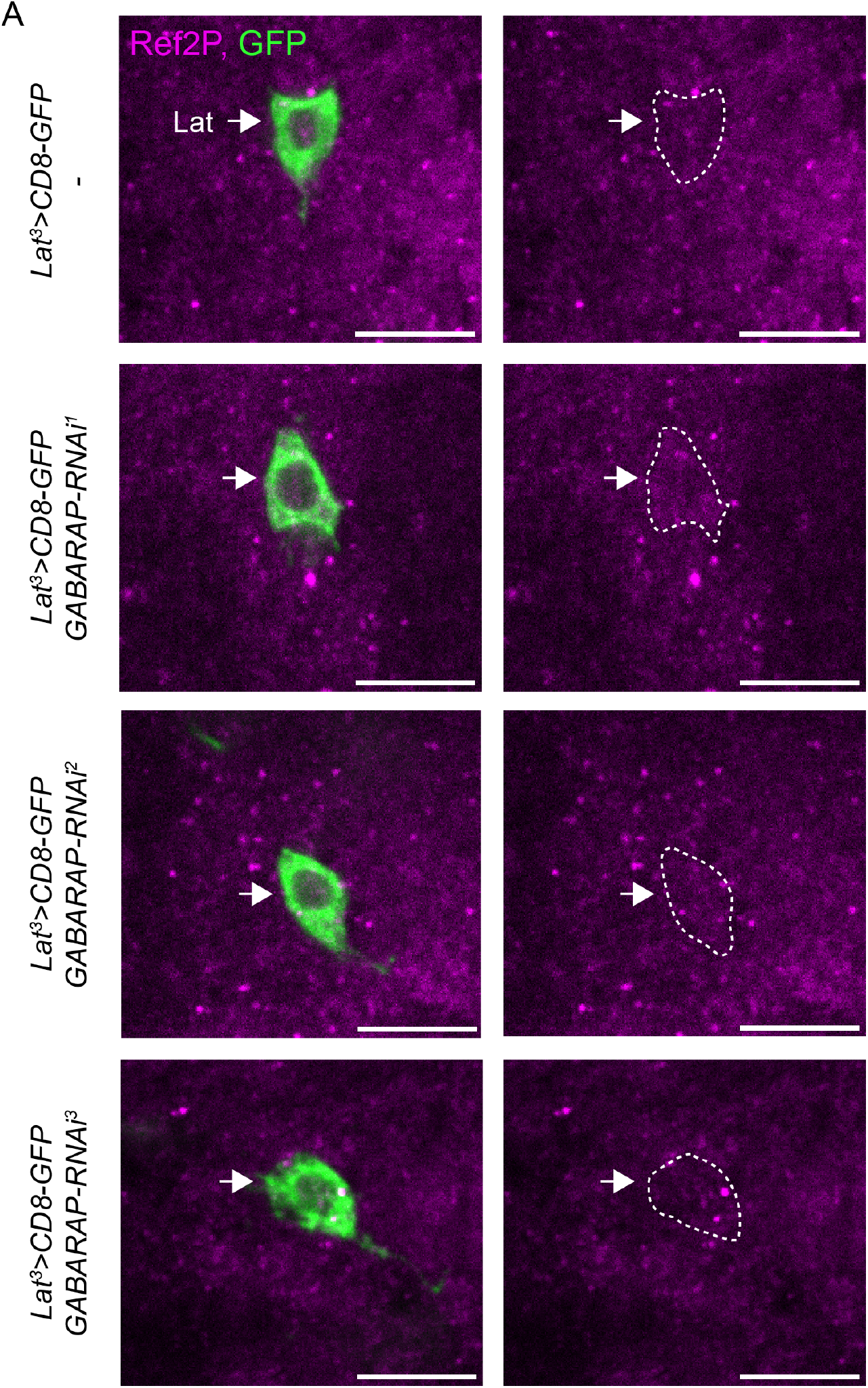
*GABARAP* knockdown in Lat neurons does not cause Ref2P accumulation, related to Figure 3. (A) *Lat*^*3*^*>CD8-GFP* male brains are stained with anti-GFP (green) and anti-Ref2P antibodies (magenta). The left column shows the labeling of a Lat cell body (green) and Ref2P protein expression (magenta) in control flies and flies expressing different *GABARAP-RNAi* hairpins. The right column shows the same image on the left without GFP staining; the location of the Lat cell body is highlighted and indicated by a white arrow (scale bars=10μm, single plane image).

**Figure S4.**
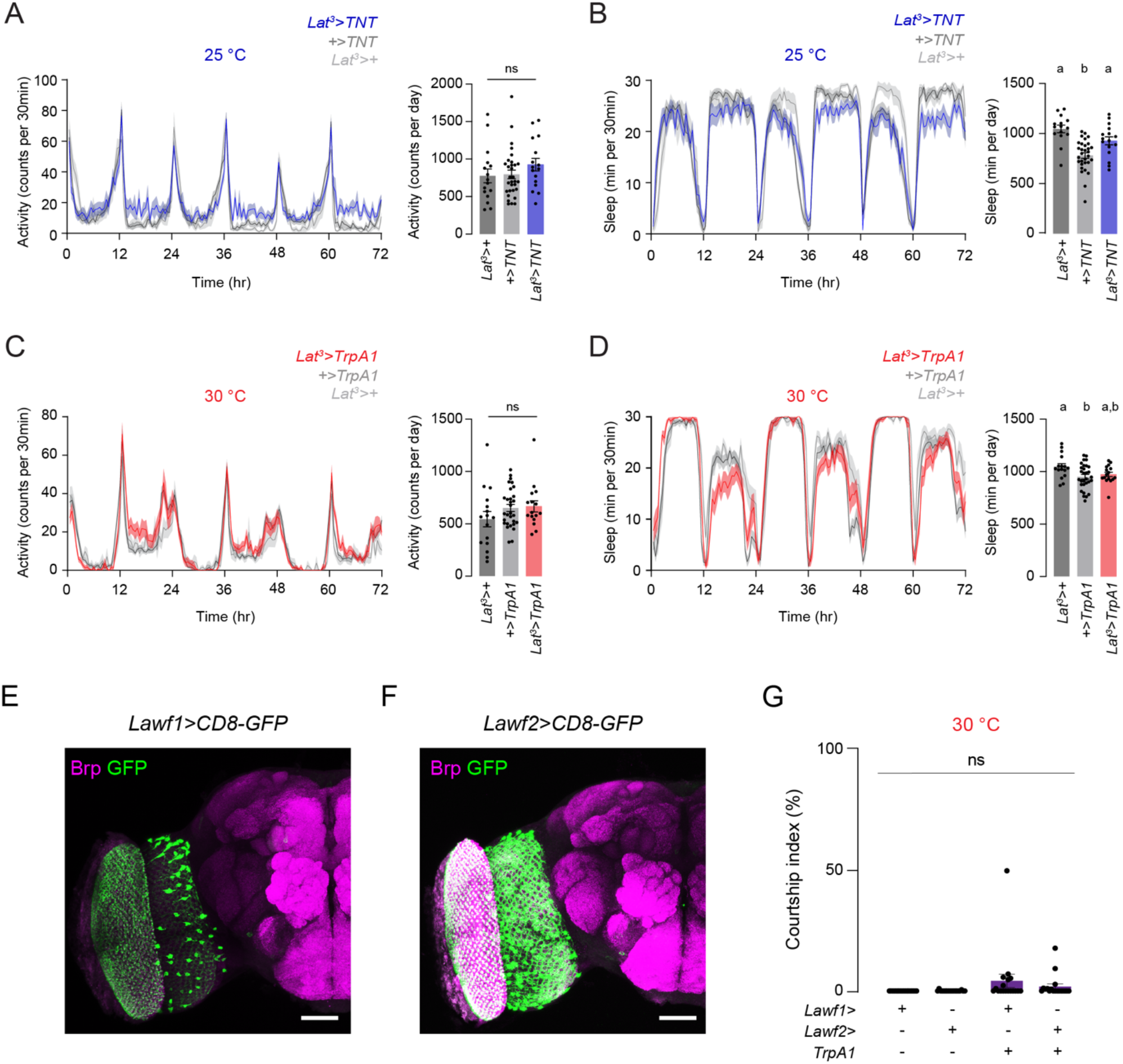
Inhibition of Lat neurons does not affect locomotor activity or sleep, and activation of other visual feedback neurons does not induce male-male courtship, related to Figure 4. (A) Circadian activity profiles of *Lat*^*3*^*>TNT* and control males plotted over 72 hours (left) or averaged per day (right) (One-way ANOVA with Tukey’s test, ns, mean ± SEM, n=16-31 flies). (B) Sleep profiles of *Lat*^*3*^*>TNT* and control males plotted over 72 hours (left) or averaged per day (right) (One-way ANOVA with Tukey’s test, p<0.05, mean ± SEM, n=16-31 flies). (C) Circadian activity profiles of *Lat*^*3*^*>TrpA1* and control males plotted over 72 hours (left) or averaged per day (right) (One-way ANOVA with Tukey’s test, ns, mean ± SEM, n=16-30 flies). (D) Sleep profiles of *Lat*^*3*^*>TrpA1* and control males plotted over 72 hours (left) or averaged per day (right) (One-way ANOVA with Tukey’s test, p<0.05, mean ± SEM, n=16-30 flies). (E-F) *Lawf1>CD8-GFP* and *Lawf2>CD8-GFP* male brains stained with anti-GFP (green) and anti-Brp (magenta) (scale bars=50μm). (G) Courtship quantification of *Lawf1>TrpA1, Lawf2>TrpA1* and control males toward male targets (One-way ANOVA, ns, mean ± SEM, n=18 flies).

**Figure S5.**
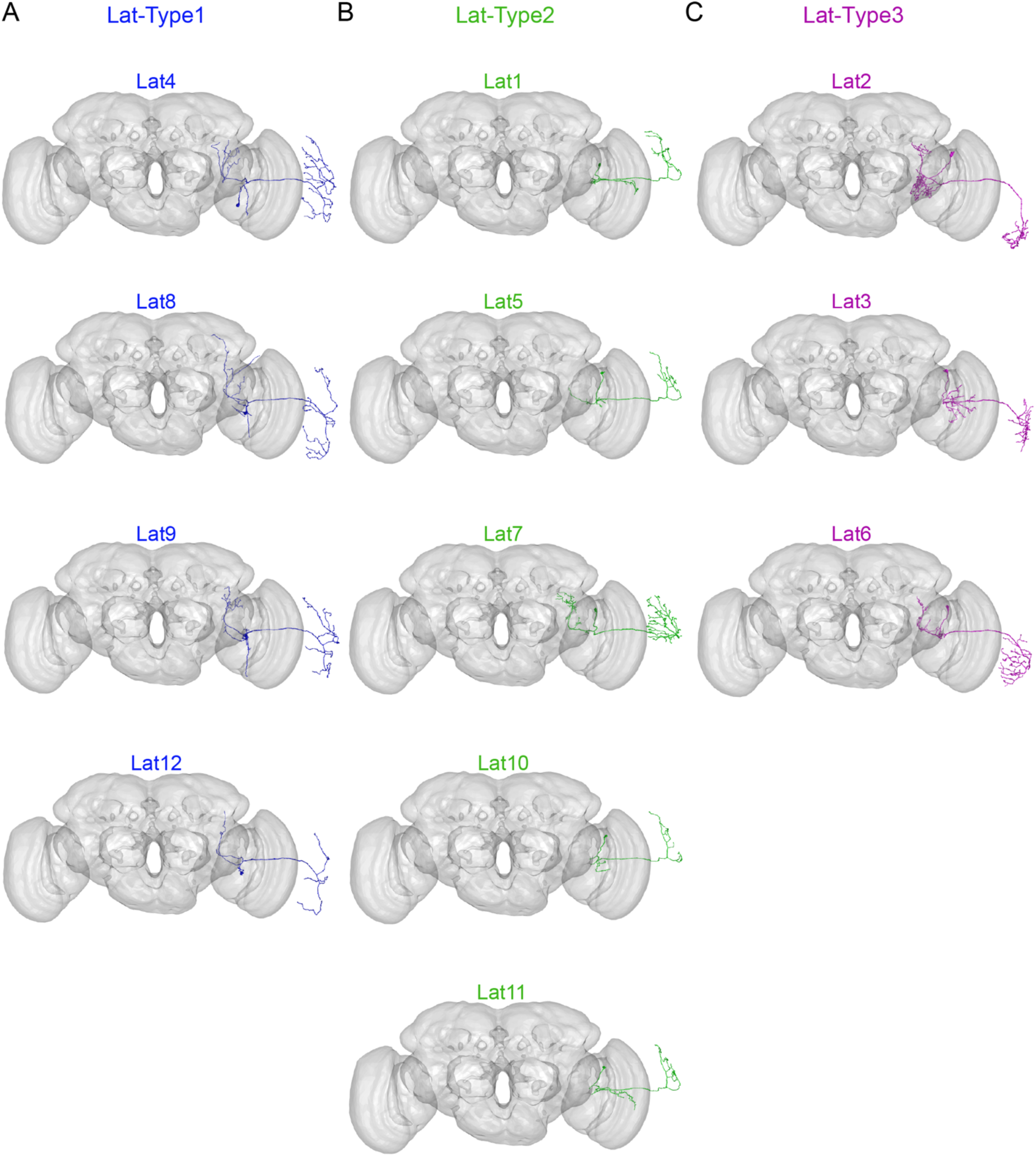
EM reconstruction of putative Lat neuron subtypes, related to Figure 6. (A) Reconstruction of putative Lat-Type1 neurons (blue, n=4) plotted against a standard FAFB14 brain mesh (gray). Lat-Type1 neurons have synaptic sites in both the dorsal and the ventral lamina. (B) Reconstruction of putative Lat-Type2 neurons (green, n=5) plotted against a standard FAFB14 brain mesh (gray). Lat-Type2 neurons have synaptic sites only in the dorsal lamina. (C) Reconstruction of putative Lat-Type3 neurons (magenta, n=3) plotted against a standard FAFB14 brain mesh (gray). Lat-Type3 neurons have synaptic sites only in the ventral lamina.

**Figure S6.**
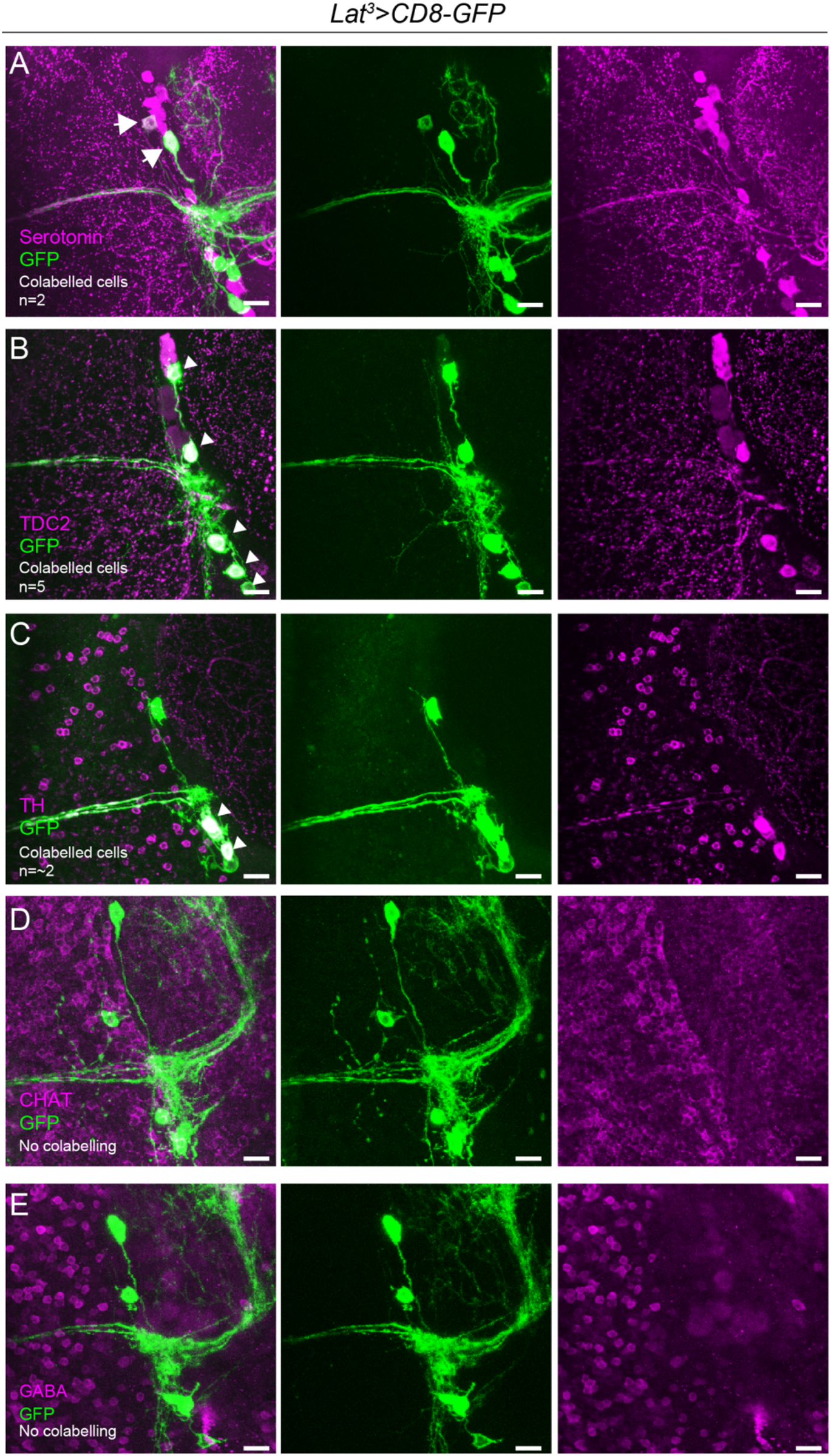
Lat neurons express multiple neurotransmitters, related to Figure 6. (A-E) *Lat*^*3*^*>CD8-GFP* male brains are stained with anti-GFP (green) and various neurotransmitter antibodies (magenta) (scale bars=10μm): (A) anti-serotonin, (B) anti-TDC2, (C) anti-TH, (D) anti-ChAT, (E), and anti-GABA. White arrowheads indicate cell bodies of Lat neurons. The number of Lat neurons labeled by each neurotransmitter is shown in the images in the left column.

**Figure S7.**
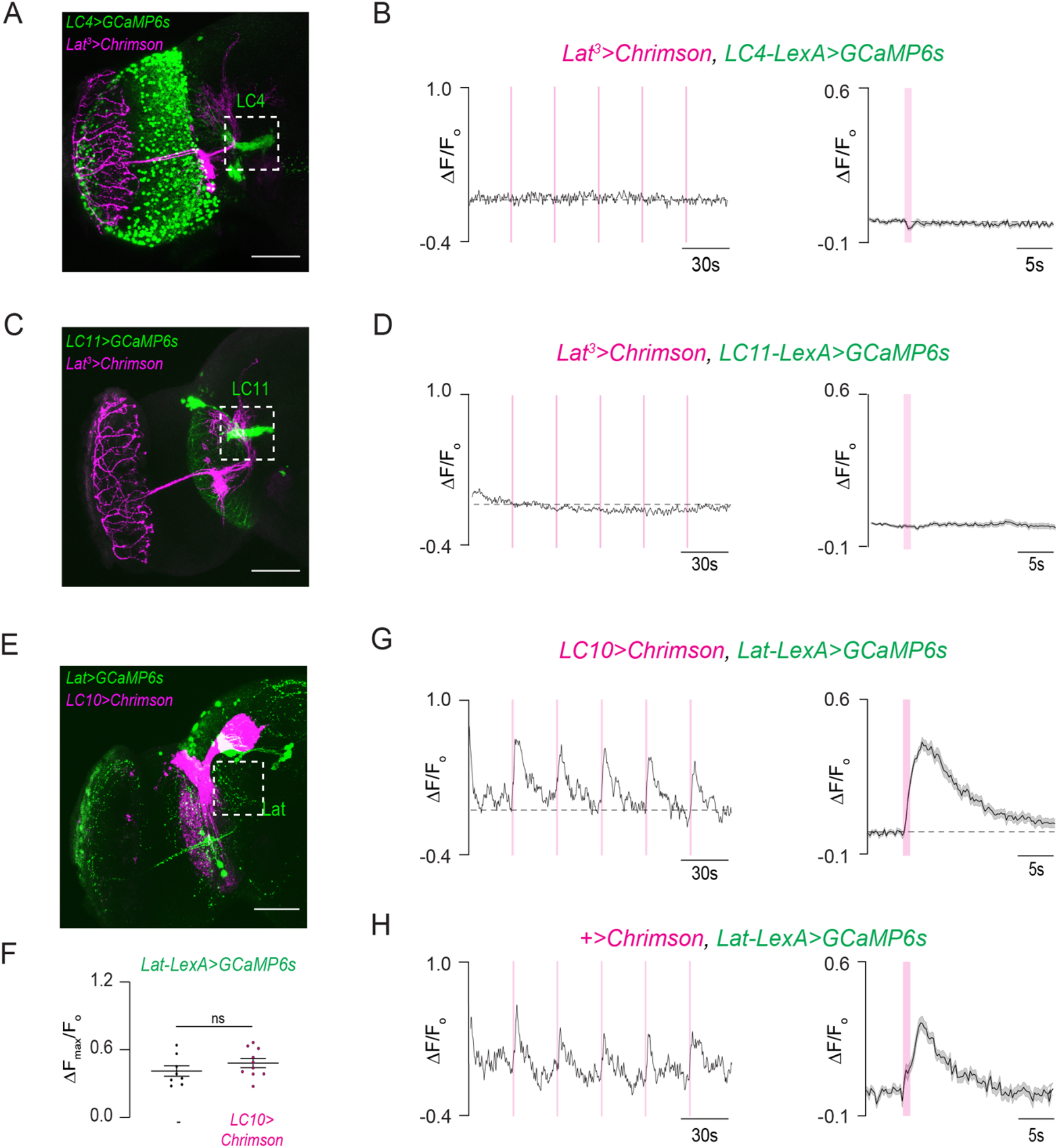
Optogenetic stimulation of Lat neurons does not activate LC4 or LC11 visual projection neurons, nor stimulation of LC10a neurons activates Lat neurons, related to Figure 7. *(A) Lat*^*3*^*>Chrimson* (magenta), *LC4-LexA>GCaMP6s* (green) example brain (scale bar= 50μm). (B) Individual (left) and averaged (right) normalized responses of LC4 neurons upon optogenetic stimulation of Lat neurons (mean ± SEM; n=10 flies). *(C) Lat*^*3*^*>Chrimson* (magenta), *LC11-LexA>GCaMP6s* (green) example brain (scale bar= 50μm). (D) Individual (left) and averaged (right) normalized responses of LC11 neurons upon optogenetic stimulation of Lat neurons (mean ± SEM; n=10 flies). *(E) LC10>Chrimson* (magenta), *Lat-LexA>GCaMP6s* (green) example brain (scale bar= 50 μm). (F) Peak normalized responses of Lat neurons upon optogenetic stimulation of LC10 neurons (Unpaired t-test with Welch’s correction; ns; mean ± SEM, n=8-10 flies). (G-H) Individual (left) and averaged (right) normalized responses of Lat neurons in *LC10>Chrimson* and control flies upon optogenetic stimulation (mean ± SEM; n=8-10 flies).

## MATERIALS AND METHODS

### EXPERIMENTAL MODEL AND SUBJECT DETAILS

#### Flies

All flies *(Drosophila melanogaster)* were maintained on conventional cornmeal-agar-molasses medium at ∼25°C, ∼65% relative humidity, and under a 12-hour light/dark cycle (lights on at 9 AM) throughout development and adulthood unless otherwise stated. Flies expressing TrpA1 were reared under a 12-hour light/dark cycle at 18°C and tested at 30°C. Lat split-Gal4s tested in all courtship assays carried a wild copy of the white gene on the X chromosome. Fly stocks are detailed in the Key Resources Table. The complete genotypes used in each figure are listed in Table S2.

## METHOD DETAILS

### Male-male courtship assay

All male-male courtship assays were performed at 25°C and 60% relative humidity from 3 PM to 7 PM unless otherwise stated. Experimental flies were collected as virgins, group-housed, and aged for 5-7 days. 5–7-day old virgin *w*^*1118*^ males were used as targets for experimental flies. Courtship assays were performed in acrylic chambers (10mm diameter x 5mm height) with a removable strip separating the chamber into two halves. Experimental and target males were placed in each half of the chamber and allowed to acclimate for 5min. The assay was initiated by removing the strip and allowing the flies to interact. Courtship behavior was recorded using a FLIR Blackfly-S camera (BFS-U3-31S4M-C or BFS-U3-13Y3M, Monochrome) at 30fps for 10-15min under white LED illumination. Flies used in TrpA1 neuronal activation experiments were aged at 18°C, and the courtship assays were performed at 30°C. To test male courtship behavior in the dark, flies were illuminated by an IR light in a light-tight enclosure, and the fly behavior was recorded using a FLIR Blackfly-S camera (BFS-U3-31S4M-C or BFS-U3-13Y3M, Monochrome) equipped with an IR filter (Edmund Optics, IR Pass Filter, R-72).

### Male-female courtship assay

To test male-female courtship behavior, we used 5–7-day old virgin male flies and 3–5-day old virgin female flies. Both males and females were reared in single-sex groups. The male-female courtship chamber (30mm diameter and 3mm height) was designed based on a previous report (Simon and Dickinson, 2010). The freely interacting male and female flies were illuminated by an IR light (Back-lit Backlights, BL1212, Advanced Illumination). To avoid heating, the arena was equipped with a liquid cooling system (Exos-2 Liquid Cooling System, EX2-750BK, Koolance). Courtship behavior was recorded at 30fps using a FLIR Blackfly-S camera (BFS-U3-31S4M-C, Monochrome) equipped with an IR filter (Edmund Optics, IR Pass Filter, R-72). The recordings lasted for 60 minutes or until successful copulation.

### Circadian activity and sleep measurements

We used 5–6-day old virgin male flies to quantify circadian activity and sleep. Flies were individually loaded into plastic tubes containing fly food. Activity data were collected in 1-min bins using the *Drosophila* Activity Monitors (DAM2, Trikinetics, Waltham, MA) for four days under a 12-hour light/dark cycle. DAM monitors contain a single IR beam positioned in the middle of the plastic tubes. When the fly walks back and forth, the IR beam gets interrupted, leaving a record. Total beam breaks were captured and summed per minute. Flies were allowed to acclimate to the DAM monitors for 24 hours, and the data from that day were not used for analysis. Neuronal silencing experiments were performed at 25°C, whereas thermogenetic activation experiments were performed at 30°C.

### Immunohistochemistry

Immunohistochemistry for the fly brain and ventral nerve chord was performed with minor modifications (Yapici et al., 2016). Briefly, flies were gently dipped into 70% ethanol for less than 30s to remove cuticular hydrocarbons on the fly body. For whole-mount staining, brains and ventral nerve cords were dissected in phosphate-buffered saline (PBS, calcium- and magnesium-free; Lonza BioWhittaker, #17-517Q) and incubated in 4% paraformaldehyde (PFA) in PBS for 30 min at room temperature on an orbital shaker. Tissues were washed with PBS at room temperature (4 times, 15 min each). Samples were blocked in 5% normal goat serum (NGS, Jackson Labs, 005-000-121) in PBS containing 0.2% Triton X-100 (PBT) for 1 hour and then incubated with primary antibodies diluted in NGS+PBT for 48 hours at 4°C. After incubation, samples were washed 5-6 times over 1 hour in 0.2% PBT at room temperature and incubated with secondary antibodies diluted in NGS+PBT for an additional 24 hours at 4°C. On the fourth day, samples were washed 4-6 times over 1 hour in PBS at room temperature and mounted with SlowFade™ Diamond Antifade Mountant (Thermo Fisher, S36972) on glass slides (VWR, #16004-368). The samples were covered by a glass coverslip on top and sealed using clear nail polish. For primary antibodies, we used mouse anti-Bruchpilot (1:20), mouse anti-GFP (1:100), rabbit anti-GFP (1:3000), chicken anti-GFP (1:5000), rabbit anti-DsRed (1:1000), rabbit anti-serotonin (1:2000), rabbit anti-Tdc2 (1:200), mouse anti-TH (1:1000), mouse anti-ChAT (1:1000), rabbit anti-GABA (1:25), rabbit anti-Ref2P (1:500). Secondary antibodies used were goat anti-mouse Alexa 488 (1:1000), goat anti-rabbit Alexa 488 (1:1000), goat anti-chicken Alexa 488 (1:1000), goat anti-rabbit Alexa 546 (1:1000), and goat anti-mouse Alexa 546 (1:1000). Z-stacks of high-resolution (1024×1024 pixels) image frames were collected at 1-2 μm intervals using an upright confocal microscope (Zeiss LSM880). Abbreviations used for serotonergic neural populations are based on a previous publication (Pooryasin and Fiala, 2015): ALP, anterior lateral protocerebrum; AMP, anterior medial protocerebrum; ADMP, anterior dorsomedial protocerebrum; LP, lateral protocerebrum; SEL, lateral subesophageal ganglion; SEM, medial subesophageal ganglion. PLP, posterior lateral protocerebrum; PMPD, posterior medial protocerebrum, dorsal; PMPM, Posterior medial protocerebrum, medial; PMPV, Posterior medial protocerebrum, ventral.

### Fly preparation for optogenetics

5–7-day old virgin males were used in all optogenetics/functional imaging experiments. Virgin males were collected after eclosion and aged for five days on a conventional cornmeal-agar-molasses medium at 25°C. Flies were transferred to a vial containing a mixture of fly food and 400μM all-trans-retinal (ATR, Sigma # R2500). Flies were kept in the dark at 25°C for an additional 1-2 days before getting tested in the optogenetics/functional imaging experiments.

### Fly preparation for two-photon imaging

Flies were anesthetized on ice and tethered to a custom fly holder designed based on a previous publication (Seelig et al., 2010). We fixed the fly’s head to the holder using a UV-curable adhesive (Bondic UV resin, #SK8024). Flies were allowed to recover from cold anesthesia in a dark incubator set to 25°C for one to two hours. Once the fly’s head was fixed, the tethering plate was filled with physiological saline (108mM NaCl, 5mM KCl, 8.2mM MgCl_2_·6H_2_O, 2mM CaCl_2_·2H_2_O, 4mM NaHCO_3_, 1mM NaH_2_PO_4_, 5mM Trehalose·2H_2_O, 10mM Sucrose, 5mM HEPES, pH 7.5). The cuticle behind the head between the eyes was dissected to gain optical access to the fly brain. The head cuticle was cut with a 16-gauge needle, and the air sacks, fat bodies, and trachea around the imaging window were removed using forceps. During all functional imaging experiments, flies stood or walked on an air-suspended, spherical treadmill (Aragon et al., 2022). Fly behavior was captured at 30 fps under IR illumination using two cameras (FLIR Blackfly-S, BFS-U3-13Y3C, and BFS-U3-16S2M) and the SpinView software (FLIR).

### Two-photon calcium imaging coupled with optogenetic stimulation

All functional imaging experiments were performed using a resonant scanning two-photon microscope (Bergamo II, Thorlabs) equipped with a 16X Plan Fluor Objective (Nikon, N16XLWD-PF). We used a Ti: Sapphire laser (Coherent Chameleon Vision II) centered at 920 nm as the two-photon excitation source. The laser was directed through a resonant scanning galvanometer for fast-scanning volumetric imaging, and a piezo-electric Z-focus controlled the objective. Volumetric imaging was achieved by scanning through 8 z-planes separated by 5μm at a volumetric scanning rate of ∼4.6Hz and 256×256-pixel resolution. Laser power was measured using a power meter (PM100D with S175C, Thorlabs). Laser power after the objective ranged between ∼12-25mW based on the expression level of GCaMP. Optogenetic stimulation was achieved using a 617nm LED that has been integrated into the light path of the two-photon microscope, and the LED light was delivered to the fly brain through the objective. LED light intensity (0.25-0.28mW) was measured by an optical power meter (PM100D with S405C, Thorlabs) placed under the objective. A long-pass filter (FELH0600, Thorlabs) was used to reduce the background elevation caused by 617nm-LED stimulation used for optogenetic stimulation. All optogenetic activation experiments started with 30s scanning without stimulation to capture baseline GCaMP fluorescence. Next, LED light stimulation was delivered to the fly brain for 1s (20Hz, 25ms ON, 25ms OFF). The stimulation was repeated five times at 30s intervals. At the end of each imaging experiment, we assessed the flies’ health condition by mechanical stimulation of the leg using forceps. The data collected from flies that did not respond to the mechanical stimulus were excluded from the final data analysis because we considered those flies unhealthy. Flies that showed substantial movement in the Z-direction were also discarded on rare occasions because of the severe motion artifacts in the imaging data.

## QUANTIFICATION AND STATISTICAL ANALYSIS

### EM reconstructions

We used the Flywire open-source platform to identify and classify the putative Lat neurons (Dorkenwald et al., 2022). We first identified putative Lat neurons based on light microscopy data, projection patterns, and cell body locations. Next, identified Lat neuron candidates were proofread using the tools provided by the Flywire platform, mainly focusing on removing incorrect annotations and focusing on the backbone of a particular neuron. We only identified Lat neurons in the left hemisphere of the whole female brain EM volume (Zheng et al., 2018) since the lamina on the right hemisphere was significantly damaged. To further analyze and plot the Lat neurons in a standard brain mesh, we used the natverse package in R-studio, a toolbox for combining and analyzing neuroanatomical data (Bates et al., 2020). Natverse consists of multiple R-packages that allow the analysis of light microscopy and EM datasets across various model organisms, including *Drosophila melanogaster*. We mainly used the R “fafbseg” package to access and analyze the Flywire datasets. Details of the “fafbseg” package can be found at https://natverse.org/fafbseg/. First, we downloaded neuron meshes for each putative Lat neuron from Flywire into the R environment and visualized them in 3D using the FAFB14 standard brain mesh. Next, we used Flywire to automatically detect synaptic sites across putative Lat neurons (n=12) and filtered out synapses with a cleft score threshold of 50 to remove false positives using the flywire_adjacency_matrix function. Finally, the connectivity matrix was generated using the synaptic connectivity data, and heatmaps were generated in R using the “pheatmap” package for visualization.

### Male-male courtship analysis

Male-male courtship videos were scored manually. We quantified the total duration in which the test male displayed any courtship behaviors (orientation, tapping, chasing, licking, singing, or abdominal bending) toward the target male. The courtship index (CI) was calculated using the following formula: CI= (Time courting/10min) x100.

### Male-female courtship analysis

Male-female courtship videos were quantified using the multi-animal pose tracking software SLEAP (v1.1.5) (Pereira et al., 2019; Pereira et al., 2022). To train the SLEAP neural network, we used the multi-animal bottom-up training model. 14 nodes were selected to track specific body parts (eyes, head, thorax, wing tips, and abdominal tip) of freely interacting pairs of virgin male and female flies. After the training, the network tracking accuracy was assessed and corrected until each body part was tracked accurately in the training dataset. The final training data set contained 683 labeled frames across 35 male-female courtship videos. The trained model was used to track all male-female fly courtship videos. All tracked videos were proofread manually, and corrections were made if the model swapped the identity of the male and female flies on rare occasions. Using the tracking data, we used a custom-written code in Python to quantify and plot male courtship steps toward the female fly. The following parameters were used to determine each courtship step as previously described with minor changes (Ribeiro et al., 2018).

1. *<u>Distance</u>*: The absolute distance between the male and female fly during courtship was calculated using the XY positions of the male and female thorax. The probability density function of the male-female distance was plotted per genotype with the following criteria: female speed >5mm/s
2. *<u>Heading angle</u>*: The male heading angle was defined as the angle between the midline of the male body and the line connecting the male and female thoraxes. The probability density function of the male heading angle (θ) was plotted per genotype with the following criteria: female speed >5mm/s
3. *<u>Following</u>*: Criteria used to define the male following: Distance between the male and female fly: 2-5mm, male heading angle (θ): <30°, walking speed of males: >1mm/s, walking speed of females:>5mm/s. All parameters should persist longer than 0.3s.
4. *<u>Stationary Orientation</u>*: Criteria used to define the male orientating: male heading angle (θ): IθI<60°, male walking speed: <1mm/s. All parameters should persist longer than 1s.
5. *<u>Wing extension</u>*: The wing angle (α) is defined as the angle of wings formed by the middle line of the male and the line connecting the wing tips to the thorax. Criteria used to define the male wing extension: Male wing angle (α): >30° for more than 0.5s.

Using the courtship steps described above, male courtship ethograms were generated. We used a hierarchical structure (orienting >following >wing extension >copulation) to plot ethograms because male flies occasionally showed simultaneous courtship actions (e.g., following and wing extension) during courtship. Each behavior index (wing extension, following) was calculated by dividing the total number of frames labeled by that courtship step by the total number of frames until copulation or the entire video length (18,000 frames at most).

### Analysis of circadian activity and sleep

Circadian activity and sleep data were analyzed using the Sleep and Circadian Analysis MATLAB Program (SCAMP) (Donelson et al., 2012). To plot daily activity and sleep data, the number of beam breaks and sleep time were binned every 30 min and plotted across time over 72 hours. A sleep bout was defined as inactivity lasting at least five minutes, based on standards in the field (Qian et al., 2017; Shaw et al., 2000). We compared the average activity and sleep duration per day across different genotypes.

### Calcium Imaging Data Analysis

Volumetric imaging stacks were first mean-projected in the z-axis and then registered using a custom-written MATLAB code. ROI selection, average fluorescence computation, ΔF/F_0_ calculation, and plotting were completed using custom scripts written in MATLAB or Python. To obtain ΔF, the background signal and the baseline fluorescence (F_0_=average of frames 5s before stimulus onset (t=-1s to -6s)) were subtracted from the raw GCaMP fluorescence time series. The difference (ΔF) was then divided by F_0_ to obtain ΔF/F_0_ (ΔF/ F_0_ = (F_(t)_ - F_0_)/F_0_). The data were aligned to the optogenetic stimulation (t=0 is when the 617nm LED is turned on). ΔF/F_0_ values were plotted against time. For statistical analysis, we analyzed the peak ΔF/F_0_ within 5s (t=0s to 5s) after each optogenetic stimulus. Each fly received 5X optogenetic stimulation, and the peak responses were averaged per fly for statical comparisons. The sample size reported in imaging experiments represents the number of flies tested. For all imaging experiments, a region of interest (ROI) was manually selected as indicated below:

| *Functional connectivity experiment* | *ROI* |
| --- | --- |
| *Lat>>LC10a* | *LC10a axon bundles close to the AOTu.* |
| *Lat>>LC4* | *LC4 axon terminals in PVLP* |
| *Lat>>LC11* | *LC11 axon terminals in the PVLP* |
| *LC10>>Lat* | *Lat dendritic arbors* |
| *LC10a>>Fru-P1* | *Lateral protocerebrum complex* |

## Statistical Analysis

All statistical analyses were performed using Prism V9.3.1 (GraphPad). Details of statistical methods are reported in the Figure legends. In all figures, plots labeled with different letters are significantly different from each other.

